# Spatial profiling of Kawasaki disease cardiac inflammation identifies STING as a therapeutic target

**DOI:** 10.64898/2026.10.16.757759

**Authors:** Benjamin L. Ross, Asli E. Atici, Youngho Lee, Paul Ngai, Emily Schahrer, Malcolm E. Lane, Michael F. Fishbein, Yuki Yokouchi, Kei Takahashi, Moshe Arditi, David Zemmour, Magali Noval Rivas

## Abstract

The cellular subsets and *in situ* mechanisms underlying the development of coronary arteritis in Kawasaki disease (KD), the leading cause of acquired heart disease in children, remain poorly defined. We characterized the cellular composition and spatial organization of cardiac inflammation in autopsy coronary artery (CA) specimens from KD patients using 64-plex immunofluorescence. We identified spatially organized immune infiltrates of neutrophils, macrophages, and T cells associated with arterial wall destruction and remodeling, as well as disrupted stromal–muscle cell colocalization and diffuse myocardial inflammation. Spatial analysis also revealed expression of the innate immune sensor STING in myeloid cells in KD CA tissues. Pharmacological and genetic inhibition of STING reduced vasculitis development in a mouse model of KD. These findings define the spatial immune architecture of KD cardiac inflammation, reveal inflammatory injury beyond the coronary arteries, and identify STING as a potential therapeutic target for treatment-resistant or severe KD.

## INTRODUCTION

Kawasaki disease (KD) is a pediatric febrile vasculitis of unknown etiology affecting small and medium-sized vessels and is the leading cause of acquired heart disease in children worldwide ^1,2^. If untreated, KD leads to systemic arteritis and the development of coronary artery (CA) aneurysms in up to 25% of cases ^2^. Timely treatment with intravenous immunoglobulin G (IVIG) reduces mortality to <0.1% and decreases the incidence of CA aneurysms to approximately 4% ^2^. However, up to 20% of KD patients do not respond to the initial treatment and are at increased risk of cardiovascular complications, such as CA aneurysms, thrombosis, stenosis, and myocardial infarction ^2–4^. KD is an immune-mediated vasculitis, associated with elevated levels of circulating pro-inflammatory cytokines IL-1β, TNF-α, IL-18, and IL-6, and local inflammatory infiltrates ^5–7^. However, the specific *in situ* mechanisms driving this hyperinflammatory response and their contribution to disease progression remain incompletely understood, limiting the development of targeted therapeutic strategies for KD.

Experimental murine models of KD, such as the *Lactobacillus casei* cell wall extract (LCWE) model of vasculitis, recapitulate several pathological and immunological features of the human disease, including coronary arteritis, myocarditis, aneurysm formation, luminal myofibroblast proliferation, fibrosis, and later stenotic remodeling ^8–11^. This murine model has provided valuable mechanistic and pathological insights, showing the role of TLR2/MyD88 sensing, NLRP3– caspase-1 activation, IL-1 signaling, and recruitment of myeloid and T cells into coronary and aortic lesions ^8,9,12,13^. However, direct investigation of affected human CA tissue from KD patients remains essential to define the *in situ* cellular landscape of KD CA lesions. Such studies are challenging due to the low KD mortality rate (<0.1%) ^2^ and limited access to CA tissues. Consequently, most studies rely on bulk or single-cell RNA sequencing (scRNA-seq) of whole blood or peripheral blood mononuclear cells (PBMCs), limiting insights into tissue-resident immune responses ^14–19^. A limited number of histological studies of CA tissues from KD patients have demonstrated a complex immune cell landscape, characterized by infiltration of neutrophils, macrophages, eosinophils, activated dendritic cells (DCs), T cells, and IgA^+^ plasmablasts within the CA ^5,6,20–24^.

Here, we used a spatially multiplexed protein profiling approach at single-cell resolution to define the cellular landscape of rare CA tissues from KD and control (CTRL) patients. We identified a marked remodeling of the spatial organization, characterized by the expansion of immune-rich neighborhoods in KD CA tissues and infiltrations of macrophages, DCs, neutrophils, and T cells surrounding the inflamed CA and the myocardium. Our multiplexed spatial analysis identified STING^+^ cells exclusively in CA tissues from KD patients, highlighting an immune feature specific to CA tissues. However, gene set enrichment analysis (GSEA) did not detect significant enrichment of STING-related genes in whole blood from acute KD patients, suggesting a specific role of this pathway in CA tissues from KD patients. Furthermore, we observed increased expression of STING and cGAS-STING pathway-related proteins in vascular tissues from LCWE-injected mice developing vasculitis. Genetic and pharmacological inhibition of STING significantly attenuated LCWE-induced vasculitis severity in mice. Our findings provide a spatially resolved cellular landscape for understanding KD pathogenesis and identify the STING pathway as a potential therapeutic target for patients with KD.

## RESULTS

### Cardiac inflammation in KD is spatially heterogeneous and extends beyond the CA

The widespread use of IVIG has markedly reduced acute mortality in KD, which ranges from 0.015% in Japan to 0.17% in the USA ^2^, limiting access to CA tissues for direct study of KD *in situ* pathogenesis. To address this, we analyzed archived rare CA tissue sections collected from 3 fatal KD cases (KD1 to KD3), obtained 18 to 33 days after disease onset (**Supplementary Table 1**). In patient KD1, we analyzed two anatomical CA regions (KD1a and KD1b). This patient died 33 days after fever initiation and developed aneurysms in both the left circumflex (LCx; KD1a) and right CA (RCA; KD1b; **Extended Data Fig. 1a, b**). Patient KD2 exhibited the most fulminant disease course, dying 18 days after disease onset, with a large aneurysm in the RCA, measuring 8 mm in diameter (KD2; **Extended Data Fig. 1c** and **Supplementary Table 1**). Patient KD3 passed away 23 days after disease onset and developed an aneurysm at an unknown site, potentially the left CA (**Extended Data Fig. 1d** and **Supplementary Table 1**). Histological examination using hematoxylin and eosin (H&E) and Verhoeff-van Gieson (EVG) revealed that all KD cases exhibited CA aneurysms, with thrombi observed in KD2 and KD3 (**Extended Data Fig. 1a–d)**. These lesions were associated with transmural infiltration of immune cells, morphologically consistent with myeloid and lymphoid populations. Structural alterations included fibrocellular intimal thickening, fragmentation of the elastic laminae, medial thinning, and disruption of vessel wall integrity (**Extended Data Fig. 1** and **Supplementary Table 1**). Inflammation was not restricted to the CAs, and in all KD cases, mild interstitial and perivascular immune infiltrates were also observed in the pericardium, epicardium, and myocardium with a more diffuse distribution (**Supplementary Table 1**). For comparison, we analyzed CA tissues from two control patients (CTRL1 and CTRL2), who died from causes unrelated to KD (**Extended Data Fig. 1e, f, and Supplementary Table 1**). In both CTRL CA tissues, the CA was structurally intact, with preserved elastic laminae and no evidence of inflammatory infiltration (**Extended Data Fig. 1e, f**). Together, this small but rare retrospective cohort of fatal KD cases demonstrates CA aneurysm formation, thrombosis, and transmural immune infiltration, highlighting spatial heterogeneity of inflammation across all cardiac layers within individual patients.

### Multiplexed immunofluorescence reveals the cellular landscape and structural remodeling of KD CA tissues

To quantitatively characterize the cellular composition and molecular pathways from CA tissues of KD and CTRL patients, we performed multiplex immunofluorescence using a 64-protein panel on the CosMx Spatial Molecular Imager (SMI, Bruker Spatial Biology). This approach enabled us to generate spatially resolved profiles of protein expression associated with immune and structural processes across cardiac tissues from CTRL and KD patients (**Fig. 1a**). The CA vessel wall, epicardium, and myocardium were readily identified and aligned with the corresponding H&E sections (**Extended Data Fig. 2a, b**). While the presence of stromal cells was detected in both CTRL and KD CA tissue sections, immune cells were mainly observed in CA tissues from KD patients (**Extended Data Fig. 2c** and **Extended Data Fig. 3a–g**). After quality control, we analyzed 957,834 cells, averaging 159,639 cells per sample. Unsupervised cell annotation was carried out in each sample before integrating the 64-plex protein expression data across all six CA tissue sections (**Supplementary Table 2**) using Harmony ^25^, from which we identified 20 distinct cell populations in the integrated samples (**Fig. 1b, c**). Projection of these clusters onto tissue sections revealed spatial distributions consistent with marker expression and tissue architecture (**Fig. 1d, e** and **Extended Data Fig. 2c**). Fibronectin^+^ Vimentin^+^ fibroblasts were broadly distributed across cardiac layers, while SMA^+^ vascular smooth muscle cells (VSMCs) localized to small and large vessels (**Fig. 1c-e** and **Extended Data Fig. 2c**). CD31^+^ endothelial cells (ECs) lined vessels but displayed fragmentation and abnormal mixing with SMA^+^ cells in inflamed KD vessels (**Fig. 1d, e**). Cardiac muscle cells (myocardial cells), characterized by low expression of immune lineage markers, were confined to the myocardium. CD56^+^ cells corresponded to nerve fascicles (**Fig. 1b–e** and **Extended Data Fig. 2c**). Twelve immune cell populations were identified and predominantly localized to the epicardium and inflamed CA area of KD tissues (**Fig. 1b–e** and **Extended Data Fig. 2c** and **Extended Data Fig. 3**). These included macrophages (CD11b^+^, CD14^+^, CD163^+/–^), neutrophils (CD15^+^), and DCs (CD4^+^, CD39^+^, IDO1^+^, HLA-DR^+^, CD11c^+^) (**Fig. 1c–e**). CD4^+^ and CD8^+^ T cells were also present, with some subsets expressing activation markers, such as granzyme A (GZMA) and CTLA4, consistent with an activated phenotype (**Fig. 1c–e**). A STING^+^ cell cluster (STING^+^, Her2^+^) was also detected in the CA vessel wall and myocardium of KD tissues, but not in CTRL tissues (**Fig. 1d** and **Extended Data Fig. 4a, b**).

**Figure 1.**
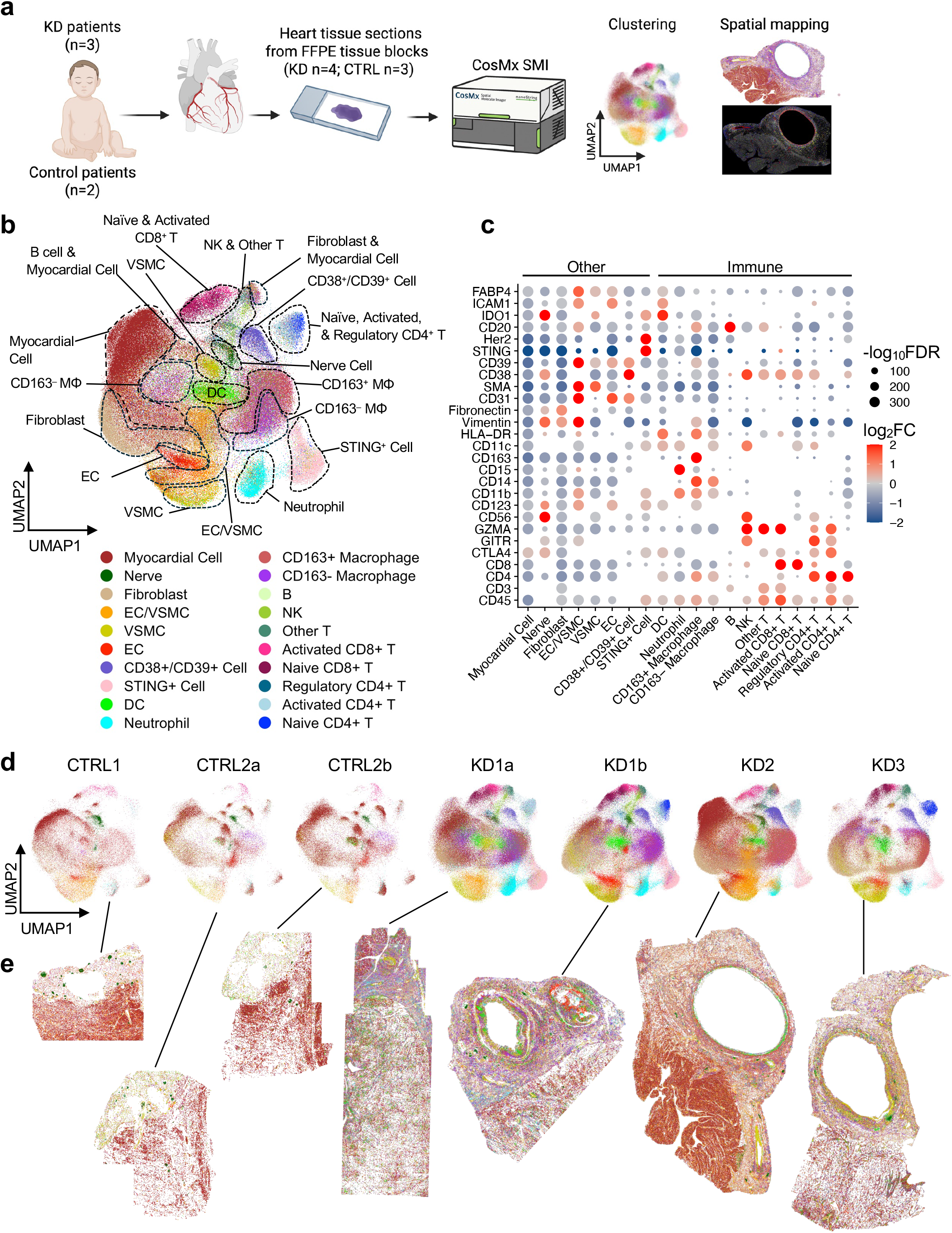
Spatial mapping of protein expression in heart tissues from controls and patients with Kawasaki disease. **a**, Schematic of the experimental design. A total of 7 coronary artery (CA) tissue sections, collected from either control patients (CTRL; n = 2; 3 sections analyzed) or patients with Kawasaki disease (KD; n = 3; 4 sections analyzed), were profiled for the spatial expression of 64 proteins using CosMx Spatial Molecular Imager (Bruker Nanostring). Schematic created with BioRender (BioRender.com). **b,** Harmonized two-dimensional (2D) UMAP embedding based on the expression of 64 proteins, highlighting 20 major cell populations across all samples. **c,** Dot plot showing expression of marker proteins across the 20 annotated cell subsets. Dot color represents the log_2_ fold change (FC; gradient of blue to red for low to high enrichment), and dot size indicates log_10_-adjusted False Discovery Rate (FDR). **d,** Individual 2D UMAP cellular embeddings of each CA tissue section highlighting the 20 major cell populations identified in each CA tissue analyzed. **e,** Spatial distribution of the identified and annotated cell types in each CA tissue. Abbreviations: CTRL, control; FFPE, Formalin-fixed paraffin-embedded; KD, Kawasaki disease; ΜΦ: macrophage; SMI, spatial molecular imager.

Multiplexed immunofluorescence enables robust *in situ* quantification and direct comparison of cell populations between KD and CTRL CA tissues (**Fig. 2** and **Extended Data Fig. 4**). In the epicardium and CA vessel wall of CTRL tissues, fibroblasts were the dominant population (519 – 739 cells/mm²), followed by resident macrophages, VSMCs, and ECs (**Fig. 2a, b** and **Extended Data Fig. 4a, b**). KD CA tissues showed a marked increase in immune cell density, particularly within the inflamed epicardium and CA wall, exceeding a 7-fold enrichment (1153.5 – 2458.8 cells/mm^2^) (**Fig. 2a, c, d** and **Extended Data Fig. 4a**). Inflammation in KD patients was not confined to the CA wall but extended diffusely across all cardiac layers (**Fig. 2c–k** and **Extended Data Fig. 4c–h**). Immune cell populations were also present in the myocardium of KD patients, with a mean total immune cell density of approximately 700 cells/mm^2^, about two-fifths the mean density of those observed in the CA vessel wall (**Fig. 2b, e, h, k**, and **Extended Data Fig. 4b, e, h**). Macrophages were markedly expanded, reaching densities above 1300 cells/mm^2^ in the vessel wall and above 400 cells/mm^2^ in the myocardium of KD patients (**Fig. 2f, h**), and included both CD163^−^ and CD163^+^ macrophages (**Extended Data Fig. 4a, b**). We observed a strong infiltration of CD163^−^ macrophages in the CA and myocardium of patient KD1, with higher densities localized to the inflamed CA vessel wall area and adjacent myocardium (**Extended Data Fig. 4a, b** and **Extended Data Fig. 5a–c**). In contrast, CA tissues from patients KD2 and KD3, both with CA thrombi (**Extended Data Fig. 1c, d**), showed lower CD163^−^ macrophage densities and increased accumulation of CD163^+^ macrophages in the epicardium, CA vessel, and myocardium (**Extended Data Fig. 4a, b** and **Extended Data Fig. 5d–f**). T cells, nearly absent in the epicardium, CA vessel wall, and myocardium of control tissues (14.07 – 101 cells/mm²), reached over 900 cells/mm² in KD CA tissues and comprised both CD4^+^, CD8^+^, and CD4^−^ CD8^−^ T cells (Other T) (**Fig. 2a, i–k** and **Extended Data Fig. 4a, b**). Neutrophils and DCs were also enriched in both the CA vessel wall and myocardium of KD patients, reflecting a coordinated innate and adaptive immune response (**Extended Data Fig. 4c–h**). In conclusion, multiplexed spatial profiling provides a quantitative framework for understanding tissue inflammation during KD, revealing a marked heterogeneity in both the cellular composition and spatial distribution of immune infiltrates that extend beyond the CA and across all cardiac tissue layers.

**Figure 2.**
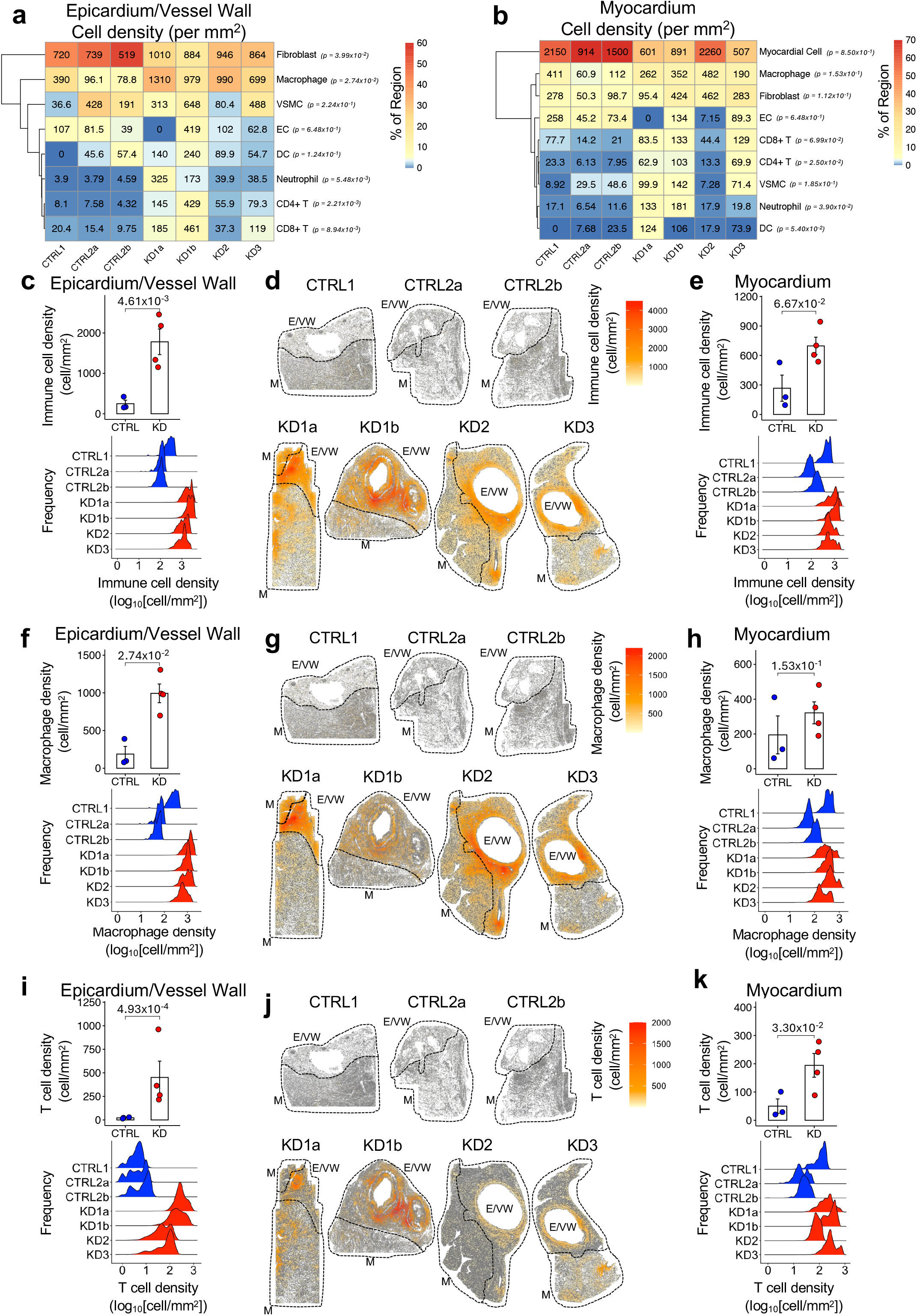
Immune cell infiltrations in coronary artery tissues of patients with Kawasaki disease. **a, b**, Heatmaps showing the density of major cell populations in the coronary artery (CA) epicardium and vessel wall (E/VW; a) and myocardium (M; b) of control (CTRL) individuals and patients with Kawasaki disease (KD). Cell density values (cell/mm^2^) are indicated within each cell. The color gradient (blue–yellow–red) represents the relative abundance (%) of each cell type for each tissue analyzed. *p*-value indicates statistical difference of the mean cell density in CTRL vs. KD samples. **c,** Bar plot and ridge plot of total immune cell density (T cells, neutrophils, dendritic cells, and macrophages) in the epicardium and CA vessel wall of CTRL and KD tissues. **d,** Spatial distribution of immune cell density in the epicardium and CA vessel wall, and myocardium of CTRL and KD tissues**. e,** Bar plot and ridge plot of total immune cell density in the myocardium of CTRL and KD tissues. **f,** Bar plot and ridge plot of macrophage density in the epicardium and CA vessel wall of CTRL and KD tissues. **g,** Spatial distribution of macrophage density in the epicardium and CA vessel wall and myocardium of CTRL and KD tissues. **h,** Bar plot and ridge plot of macrophage density in the myocardium of CTRL and KD tissues. **i,** Bar plot and ridge plot of T cell density in the epicardium and CA vessel wall of control and KD tissues. **j,** Spatial distribution of T cell density in the epicardium and CA vessel wall and myocardium of CTRL and KD tissues. **k**, Bar plot and ridge plot of T cell density in the myocardium of CTRL and KD tissues. (a-c, e, f, h, i, and k) *p-*values were calculated using a one-sided unpaired *t*-test. Abbreviations: CTRL; control, E/VW; epicardium and vessel wall, KD; Kawasaki disease, M; myocardium, VSMCs; vascular smooth muscle cells, ECs; endothelial cells, DC; dendritic cells, NK; natural killer cells.

### Spatial organization of immune infiltration reveals layered architecture and remodeling of cellular neighborhoods in KD cardiac tissues

To better understand the organization of immune infiltration within KD CA tissues, we analyzed the spatial distribution of cells, their local neighborhoods, and pairwise interaction networks ^26^. In CA tissues from CTRL patients, the cellular composition followed a well-defined spatial gradient, and ECs, VSMCs, and fibroblasts were enriched near the lumen (<400 µm) (**Extended Data Fig. 6a**). At greater distances, the cellular composition transitioned toward higher proportions of myocardial cells, consistent with the histological tissue layers and architecture of heart tissues (**Extended Data Fig. 6a**). This spatial distribution was disrupted in CA tissues from KD patients, in which higher proportions of immune cells (macrophages, T cells, neutrophils, and DCs) were observed close to the CA lumen (<400 µm) (**Extended Data Fig. 6b**). Compared to CTRL tissues, fibroblasts were also less abundant near the CA lumen (<400 µm) of KD patients, and their proportions appear to increase at further distances (>600 µm) (**Extended Data Fig. 6b**). These compositional alterations persisted at greater distances from the CA lumen (>1000 µm) of KD patients, where reduced abundance of myocardial cells, expansion of fibroblast and presence of immune cell infiltrations were observed (**Extended Data Fig. 6b**).

To determine whether CA tissue organization reflected coordinated cellular colocalization, we next analyzed cell-to-cell neighbors by constructing pairwise proximity networks based on physical distance. Comparison of CTRL and KD CA tissues revealed marked alterations in cellular colocalization, characterized by enrichment of immune cells, including macrophages, neutrophils, DCs, CD4^+^ and CD8^+^ T cells in the epicardium and CA vessel wall area of KD patients (**Fig. 3a**). In contrast, interactions among structural cell types, including fibroblasts, VSMCs, and nerve fascicles, were reduced (**Fig. 3a**). Similar changes were observed in the myocardium of KD patients, with increased colocalization between CD4^+^ T cells and neutrophils, whereas colocalization between myocardial and nerve fascicles was reduced (**Fig. 3b**). These findings indicate that immune infiltrations are associated with disruption of cellular organization across cardiac layers.

**Figure 3.**
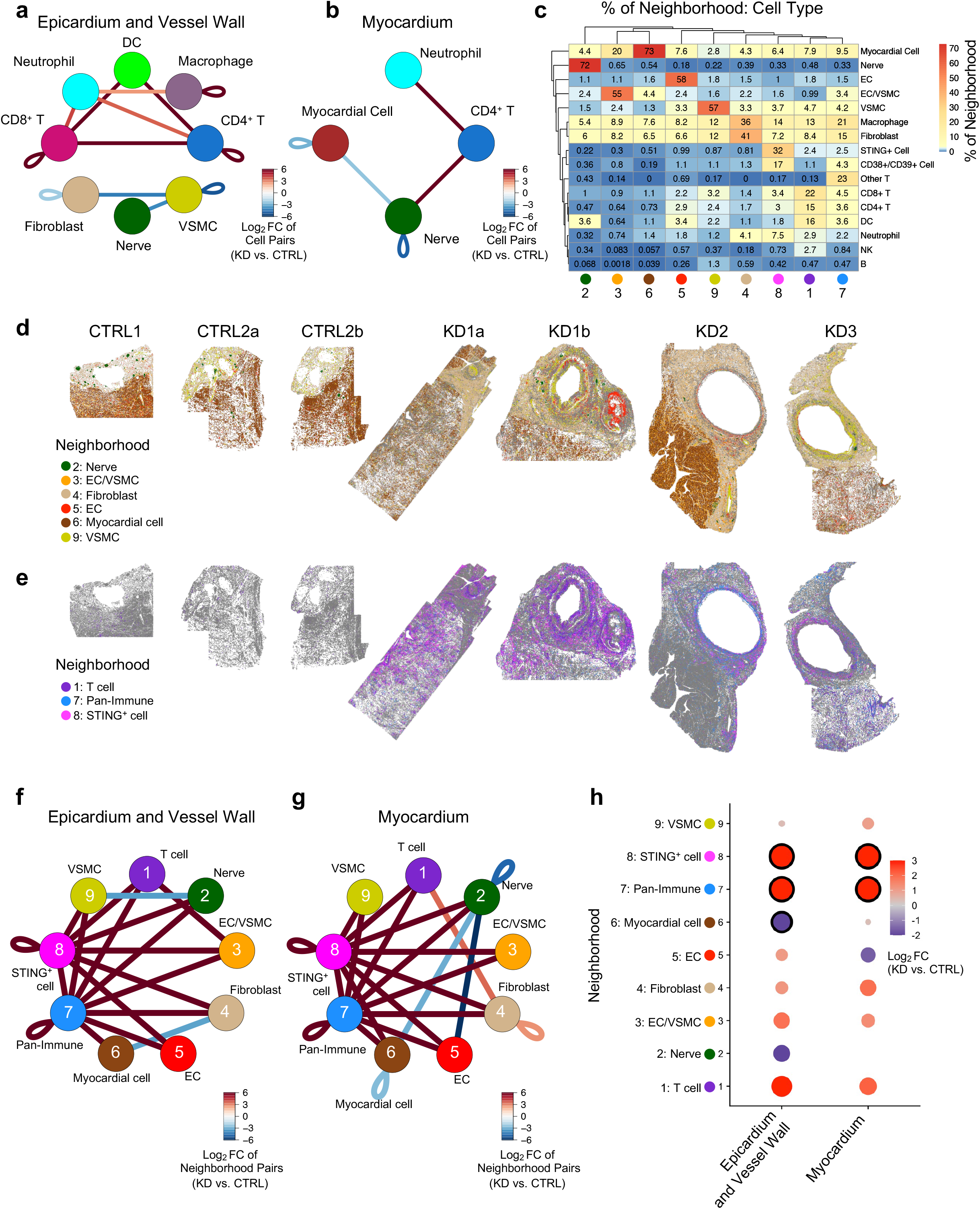
Expansion of immune-rich neighborhoods and disruption of spatial organization in coronary artery tissues from patients with Kawasaki disease. **a, b**, Network graph showing enriched colocalized cell pairs in the epicardium and coronary artery (CA) vessel wall (a) and myocardium (b) of Kawasaki disease (KD) CA tissues compared with control (CTRL). The color of the connecting lines is proportional to the log_2_ Fold Change (FC) in KD vs. CTRL CA tissues. Only statistically significant enriched cell pairs are shown (FDR <0.05, empirical Bayes moderated 2-tailed t-test). **c,** Heatmap showing the cell type composition the 9 identified neighborhoods, with a color gradient (blue–yellow–red) indicating low to high percentage of each cell type in each neighborhood. The percent of neighborhood value is indicated in each cell. **d, e,** Spatial distribution of the 9 cellular neighborhoods in each sample defined in (c). **f, g,** Network graph of enriched colocalized neighborhood pairs in the epicardium and vessel wall (f) and myocardium (g) of KD CA tissues compared with CTRL. The color of the connecting line is proportional to the log_2_ fold change (FC) in KD samples vs CTRL samples. Only statistically significant neighborhood pairs are shown (*p*-value <0.05, empirical-Bayes-moderated 2-tailed t-test). **h,** Dot plot of the log_2_ FC in cellular neighborhood density between KD vs. CTRL across either the epicardium and CA vessel wall or the myocardium. The color gradient (blue to red) represents low-to-high log_2_ FC for KD vs. control samples, with dot size proportional to the log_2_ FC value. Statistically significant enrichments *(p*-value *<*0.05, 2-tailed unpaired t-test) are outlined in black. Abbreviations: VSMCs, vascular smooth muscle cells; ECs, endothelial cells; DC, dendritic cells; NK, natural killer cells; KD, Kawasaki disease; CTRL, control.

To further generalize these observations, we defined cellular neighborhoods by analyzing the composition of each cell’s local microenvironment (k = 5 nearest neighbors) and performed unsupervised clustering of these neighborhoods (**Fig. 3c–e**). We identified 9 distinct neighborhoods across the CA vessel wall, epicardium, and myocardium in CTRL and KD CA tissue sections (**Fig. 3c–e** and **Extended Data Fig. 7a–c**). Five neighborhoods corresponded to canonical tissue structures present in both CTRL and KD CA tissues, including the intima (neighborhood 5; enriched for ECs), media (neighborhood 9; enriched for VSMCs), myocardium (neighborhood 6; enriched for myocardial cells), and nerve fascicles (neighborhood 2; enriched for nerve fascicles) (**Fig. 3c** and **Extended Data Fig. 7a, b**). A fibroblast– and macrophage-rich neighborhood (neighborhood 4) was distributed across the epicardium (**Fig. 3c, d** and **Extended Data 7c**). In addition to neighborhood 4, three other immune-enriched neighborhoods (neighborhoods 1, 7, and 8) were expanded in KD CA tissues and showed distinct cellular compositions and spatial distributions compared with those in CTRL tissues (**Fig. 3c, e** and **Extended Data Fig. 7a–c**). Neighborhood 1 was enriched in T cells, with CD8^+^ and CD4^+^ T cells accounting for 22% and 15% of its cellular composition, respectively (**Fig. 3c**). It also contains DCs (16%), macrophages (13%), and neutrophils (2.9%), and was mainly distributed in inflamed CA regions (**Fig. 3c, e** and **Extended Data Fig. 7c**). Neighborhood 8, mainly composed of STING^+^ cells (32%), CD38^+^/CD39^+^ cells (17%), and macrophages (14%) (**Fig. 3c**), was only detected in the CA tissues from KD patients, and localized close to the CA vessel wall, with some cells detected in the myocardium (**Fig. 3d, e** and **Extended Data Fig. 7a–c**).

In CTRL CA tissues, we observed different neighborhood composition with distance from the CA lumen, consistent with a layered spatial organization. Neighborhood 9 (VSMC-rich) and neighborhood 5 (EC-rich) were enriched in lumen-proximal areas (<200 µm), while neighborhood 4 (fibroblast-rich) was detected at intermediate distances from the CA lumen (200 to 600 µm) and proportions of neighborhood 6 (myocardial cell-rich) increased at further distances from the CA lumen (800 to 1000 µm) (**Fig. 3d, e** and **Extended Data Fig. 8a**). This neighborhood spatial distribution observed in CTRL tissues was less pronounced in CA tissues from KD patients. While neighborhoods 3 (EC/VSMCs-rich) and 5 (EC-rich) remained present in KD tissues, the proportion of neighborhood 6 (myocardial cell-rich), appeared reduced (**Extended Data Fig. 8b**). Furthermore, a mixture of fibroblast-rich (neighborhood 4) and immune-rich neighborhoods (1: T cell-rich, 7: pan-immune, and 8: STING^+^ cell-enriched), was observed across all distances from the CA lumen (**Fig. 3d, e, Extended Data Fig. 7c** and **Extended Data Fig. 8b**), indicating that immune-associated neighborhoods are not confined to the inflamed CA area.

We next analyzed neighborhood-to-neighborhood colocalizations by constructing pairwise proximity networks based on physical distance ^26^. Compared with CTRL tissues, the epicardium-CA vessel wall and myocardium of KD patients were enriched in colocalizing immune-rich neighborhoods, particularly neighborhoods 1 (T cell-enriched), 7 (pan-immune-enriched), and 8 (STING^+^ cell-enriched) (**Fig. 3f, g**). On the other hand, colocalization of structural and stromal neighborhood 9 (VSMC-enriched) and 2 (nerve fascicles-enriched), as well as neighborhood 4 (fibroblast-enriched) and 6 (myocardial cell-enriched), were decreased in the CA vessel wall area (**Fig. 3f**). A reduction in the density of structural neighborhoods, such as myocardial cells (neighborhood 6) and nerve fascicles (neighborhood 2), was also detected in the epicardium/CA vessel wall of KD tissue sections (**Fig. 3h**). In the myocardium, decreased cellular density in EC (neighborhood 5) was also observed in KD CA tissues (**Figure 3h**). Together, these analyses reveal that KD is characterized by the expansion of immune neighborhoods and the contraction of structural tissue compartments, accompanied by the reorganization of cellular and neighborhood interactions. This spatial remodeling highlights the coordinated, immune-mediated nature of infiltrations and their impact on tissue architecture during KD.

### Distinct spatial distribution of IL-1β^+^ cells and STING^+^ myeloid cells in CA tissues from KD patients

Consistent with previous reports indicating the systemic activation of the NLRP3 inflammasome and IL-1 pathway during KD ^13,27–29^, we detected IL-1β protein in CA tissues from KD patients, which was mainly localized to the inflamed CA area (**Fig. 4a** and **Extended Data Fig. 9**). The cellular subsets associated with IL-1β expression in CA tissues from KD patients were diverse, and included macrophages, neutrophils, fibroblasts, VSMCs, and ECs (**Fig. 4b**). STING^+^ cells were detected in the epicardium and CA vessel wall of all KD tissues, with a distribution spanning from proximal to distal areas in the CA lumen and presence in the myocardium (**Fig. 4c–e** and **Extended Data Fig. 6a, b**). Most STING^+^ cells co-localized with the myeloid markers CD45, CD11b, and CD14, and were concentrated at inflamed areas of the CA tissue (**Fig. 4d,f,g**). Considering that the IL-1β and STING pathways can overlap, we next assessed the spatial co-localization of IL-1β^+^ and STING^+^ cells in CA tissues from KD patients. STING and IL-1β only partially overlapped within the inflamed CA tissues from KD patients, being localized to adjacent, but distinct, areas (**Extended Data Fig. 10**). Together, these findings identify a STING-enriched myeloid niche within KD CA tissues which is spatially adjacent but distinct from IL-1β–producing cells, suggesting complementary, potentially cooperative inflammatory pathways.

**Figure 4.**
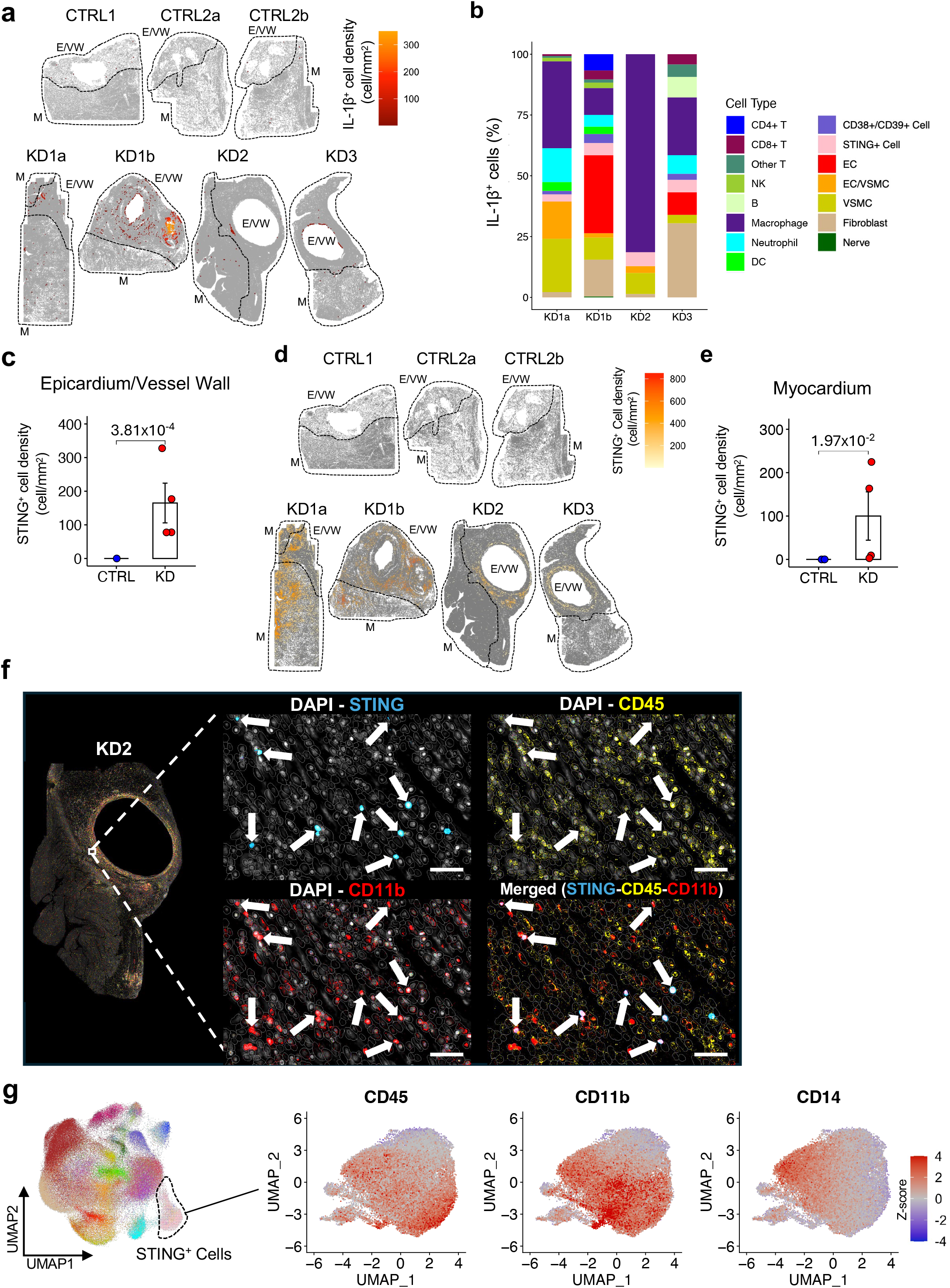
STING+ infiltrates inflamed coronary artery tissues of Kawasaki disease patients. **a**, Spatial distribution of IL-1β^+^ cells in the epicardium and coronary artery (CA) vessel wall and myocardium of patients with Kawasaki disease (KD). **b,** Bar plot showing proportions of IL-1β^+^ cells by cell type in the epicardium and CA vessel wall of KD patients. **c,** Bar plot of STING^+^ cell density in the epicardium and CA vessel wall of control (CTRL) and KD tissues. **d,** Spatial distribution of STING^+^ cells in the epicardium, CA vessel wall, and myocardium of CTRL and KD tissues. **e,** Bar plot of STING^+^ cell density in the myocardium of CTRL and KD tissues. **f,** Napari imaging of DAPI (white), STING (blue), CD45 (yellow), and CD11b (red) fluorescence in the epicardium and CA vessel wall of KD patients. White arrows indicate cells co-expressing CD45, CD11b, and STING proteins. Scale bar: 50 µm**. g,** UMAP of STING^+^ cells and scaled expression of CD45, CD11b, and CD14 among STING^+^ cells in CA tissues of KD patients. *p-values* calculated via the one-tailed unpaired t-test (c, e). Abbreviations: VSMC, vascular smooth muscle cells; EC, endothelial cells; DC, dendritic cells; NK, natural killer cells; KD, Kawasaki disease; CTRL, control; E/VW, epicardium and vessel wall; M, myocardium.

We next assessed whether transcriptional signatures associated with IL-1β and STING activation could be detected systemically during acute KD, and performed gene set enrichment analysis (GSEA) on publicly available whole blood transcriptomic datasets ^15,30^. We compared the enrichment of selected gene sets related to cGAS-STING signaling, inflammatory responses, and type I interferon responses (**Supplementary Table 3**) between patients with acute KD and healthy controls (HC), patients with acute KD and febrile controls (FC), and patients with acute KD and IVIG-treated convalescent KD patients (**Extended Data Fig. 11a–c**). GSEA detected positive enrichment of genes associated with interferon responses and NLRP3/IL-1β-related pathways in whole blood from patients with acute KD compared with HC and convalescent KD patients (**Extended Data Fig. 11a, c**). However, no significant positive enrichment in cGAS-STING pathways was observed in acute KD patients when compared with either HC, FC, or convalescent KD patients (**Extended Data Fig. 11a–c**). Therefore, the analysis of local immune responses revealed STING as an immune program specifically activated in CA tissues, not systemically.

### STING^+^ myeloid cells infiltrate the CA area in a mouse model of KD vasculitis

Our spatial analysis identified a population of STING^+^ cells uniquely present in CA tissues of KD patients (**Fig. 4c–g**). To determine whether activation of the STING pathway is recapitulated in a mouse model of KD vasculitis, we used the *Lactobacillus casei* cell wall extract (LCWE)-induced mouse model of KD ^8–10^. As previously reported, LCWE injection into WT mice leads to the development of aortitis and proximal coronary arteritis (**Fig. 5a**). Human KD is also associated with aneurysms and inflammation in large arteries, including the abdominal aorta ^23,31–33^. Similarly, LCWE injection induces inflammation and dilations of the abdominal aorta ^8^ (**Extended Data Fig. 12a**). We performed GSEA on a bulk RNA sequencing dataset (GSE141072 ^34^) generated from abdominal aortas of PBS– and LCWE-injected mice (**Extended Data Fig. 12b**). Among the gene sets significantly enriched in the abdominal aorta dilations of LCWE-injected mice, and consistent with previous studies ^13,34^, we observed a significant enrichment of gene sets associated with the NF-κB pathway, type I IFN responses, IL-1 signaling and the NLRP3 inflammasome (**Extended Data Fig. 12b**). The KEGG pathway cytosolic DNA sensing and the Ishikawa STING signaling signature were also significantly enriched in the abdominal aortas of LCWE-injected mice (**Extended Data Fig. 12b**). Analysis of a published scRNA-seq. dataset generated from abdominal aortas of PBS– and LCWE-injected mice (GSE178765 ^13^) showed *Sting1* expression across multiple cell types, including stromal populations (ECs, VSMCs, and fibroblasts), and immune cells infiltrating the inflamed abdominal aorta of LCWE-injected mice (**Extended Data Fig. 12c**).

**Figure 5.**
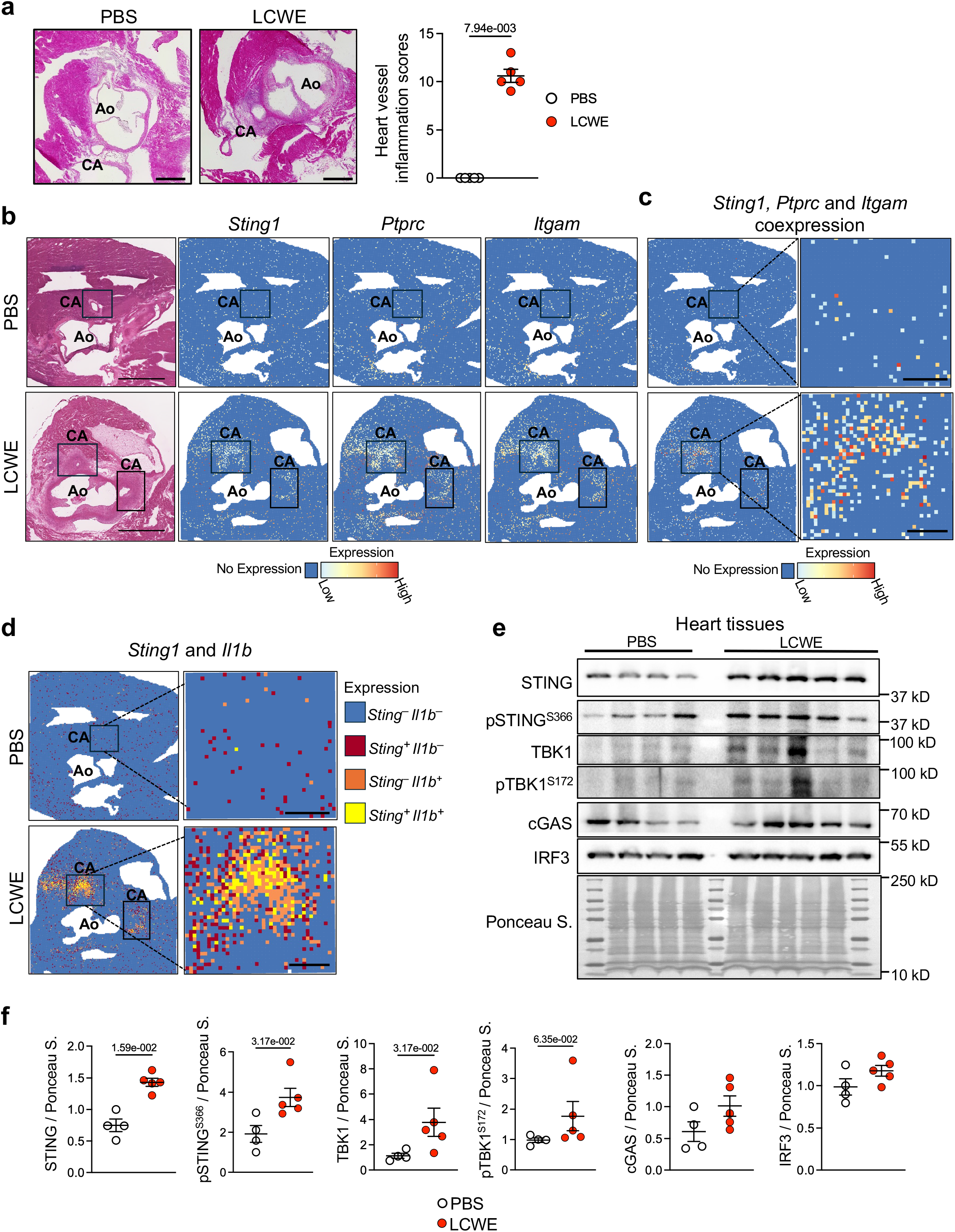
Activation of the STING pathway in heart tissues of LCWE-injected mice. **a**, H&E staining of heart tissue sections and heart vessel inflammation score of PBS– or LCWE-injected WT mice (n = 5/group). Scale bars: 500 µm. **b,** Heart tissues of PBS– and LCWE-injected WT mice were collected 2 weeks post-LCWE injection, and cryosections containing the aorta and the coronary artery (CA) were used for Visium HD transcriptomics. Annotated H&E heart tissue sections from PBS– and LCWE-injected mice, and expression of *Sting1* (STING), *Ptprc* (CD45), and *Itgam* (CD11b). Scale bars, 1000 µm. Dark blue: no expression. Light blue-yellow-red gradient: min-max normalization of log-transformed counts per million reads expression. Bin resolution: 16 µm. **c,** *Sting1*, *Ptprc*, and *Itgam* co-detection in heart tissues from PBS– and LCWE-injected mice. Inlets show a magnification of one CA area. Scale bars: 200 µm. Dark blue: no expression. Light blue-yellow-red gradient: min-max normalization of summed log-transformed counts per million reads expression. Bin resolution: 16 µm. **d,** *Sting1* and *Il1b* codetection in heart tissues from PBS– and LCWE-injected mice. Inlets show a magnification of one CA area. Scale bars: 200 µm. Dark blue: no expression. Red: *Sting1* expression. Orange: *Il1b* expression. Yellow: *Sting1* and *Il1b* co-detection. Bin resolution: 16 µm. **e,** Western blot (WB) analysis of STING, pSTING^S366^, TBK1, pTBK1^S172^, cGAS, and IRF3 in heart tissue lysates from PBS– and LCWE-injected WT mice at 2 weeks post-injection (n = 4 – 5/group). Ponceau S staining was used as the loading control. **f,** Quantification of protein expression by densitometry in heart tissue lysates from PBS– and LCWE-injected WT mice, at 2 weeks post-injection. Protein expression is presented as fold change over Ponceau S. Data are presented as mean ± SEM (a, f), with each symbol representing one individual mouse. *p-values* were obtained using the Mann-Whitney test (a, f). Abbreviations: Ao, Aorta; CA, coronary artery.

We performed high-resolution spatial transcriptomics (ST; Visium HD) analysis of heart tissues from PBS control and LCWE-injected mice and observed that *Sting1* transcripts were strongly enriched in the area surrounding the inflamed CAs (**Fig. 5b**). *Sting1* expression overlapped in these areas with *Ptprc* and *Itgam* transcripts, which encode CD45 and CD11b respectively, indicating accumulation of *Sting1-*expressing myeloid cells within the CA wall of LCWE-injected mice (**Fig. 5c**). As in patients, *Sting* and *Il1b* only partially overlapped in the tissues (**Fig. 5d**). Furthermore, compared with PBS-injected control mice, heart tissues from LCWE-injected mice showed a significant increased expression of proteins associated with *in situ* activation of the cGAS-STING pathway, including STING, phosphorylated STING (pSTING^S366^), and TBK1 (**Fig. 5e, f**). A broader increase in cGAS-STING protein expression was observed in the inflamed abdominal aorta of LCWE-injected mice, where cGAS, STING, TBK1, phosphorylated TBK1 (pTBK1^S1^^72^), and IRF3 were significantly increased compared with PBS-injected control mice (**Extended data Fig. 12d, e**). Together, these results parallel our observations in CA tissues from KD patients and support using the LCWE-induced KD model to assess the functional contribution of the STING pathway to the development of coronary arteritis.

### STING loss-of-function or pharmacological inhibition reduces the severity of LCWE-induced KD

To assess the functional contribution of the cGAS-STING pathway to LCWE-induced KD, we used Sting Goldenticket (*Sting^gt/gt^*) mice, which harbor a point mutation in *Sting* that results in a nonfunctional allele and undetectable STING protein expression ^35^. Compared to LCWE-injected WT mice, LCWE-injected *Sting^gt/gt^* mice showed decreased severity of coronary arteritis and aortitis (**Fig. 6a, b**) and reduced development of abdominal aorta dilations (**Fig. 6c, d**). LCWE injection induces an acute production of IL-1β in the peritoneal cavity ^13,36^, which was significantly reduced in LCWE-injected *Sting^gt/gt^* mice (**Fig. 6e**). LCWE-induced infiltrations of F4/80^+^ cells and Ly6G^+^ cells in heart tissues ^13,36^ were also reduced in *Sting ^gt/gt^* mice when compared with LCWE-injected WT mice (**Fig. 6f–i**). These data indicate that *Sting* loss of function protects against LCWE-induced vasculitis.

**Figure 6.**
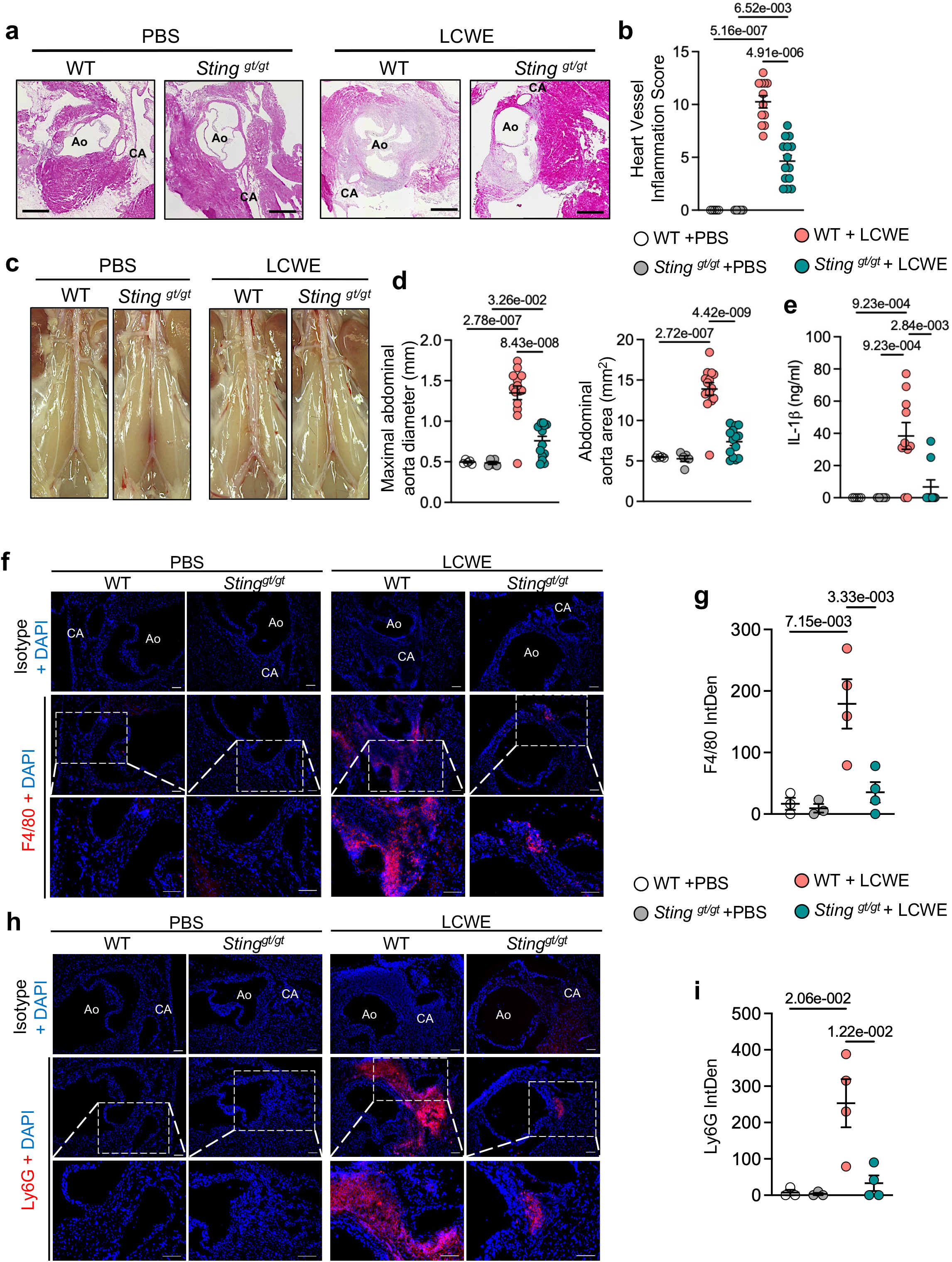
Decreased severity of LCWE-induced KD vasculitis in *Sting^gt/gt^* mice. **a, b**, H&E-stained heart sections (a) and heart vessel inflammation scores (b) of WT and *Sting^gt/gt^* mice injected with either PBS or LCWE, at 2 weeks post-injection. (n = 5-14/group). Scale bars: 500 µm. **c, d,** Pictures of the abdominal aorta area (c), maximal abdominal aorta diameter (d, left), and abdominal aorta area (d, right) measurements of WT and *Sting^gt/gt^* mice injected with either PBS or LCWE at 2 weeks post-injection (n = 5-14/group). **e,** IL-1β measurements in the peritoneal lavage of WT and *Sting^gt/gt^* mice injected with PBS or LCWE, at 24 hours post-injection (n = 5-10/group). **f, g,** Immunofluorescent staining (f) and F4/80 (red) integrated density (IntDen) quantification (g) in heart tissue from WT and *Sting^gt/gt^* mice injected with either PBS or LCWE at 2 weeks post-injection (n = 3-4/group). DAPI (blue) used to stain nuclei. Scale bars: 100 µm. **h, i,** Immunofluorescent staining (h) and quantification of Ly6G (red) integrated density (IntDen) (i) in heart tissue sections from WT and *Sting^gt/gt^* mice injected with either PBS or LCWE at 2 weeks post-injection (n = 3-4/group). DAPI (blue) used to stain nuclei. Scale bars: 100 µm. Data are presented as mean ± SEM, and pooled from 2 independent experiments (a, d). Each symbol represents one individual mouse. *p*-values obtained by 2-way ANOVA with Tukey’s multiple comparisons test (b, d, e, g, i). Abbreviations: Ao, Aorta; CA, coronary artery; IntDen, integrated density.

To determine whether pharmacological inhibition of STING signaling could reduce the severity of LCWE-induced KD vasculitis, we selected C-176 and H-151, two STING inhibitors associated with decreased systemic type I IFN production and reduced TBK1 phosphorylation in mice ^37^. WT mice were injected daily for 4 days with either C-176 or H-151 ^37^, starting the day of LCWE injection, and the severity of LCWE-induced KD vasculitis was assessed one week later (**Fig. 7a**). Both STING inhibitors reduced the development of LCWE-induced coronary arteritis and aortitis, as evidenced by a marked decrease in heart vessel inflammation (**Fig. 7b, c**) and abdominal aortic dilatation (**Fig. 7d, e**). Expression of F4/80 and STING in heart tissues of LCWE-injected mice was also reduced by STING inhibition (**Fig. 7f, g**). Altogether, our genetic and pharmacologic evidence indicates that STING blockade attenuates LCWE-induced KD vasculitis, and identifies the cGAS-STING pathway as a promising therapeutic target to mitigate KD development and progression.

**Figure 7.**
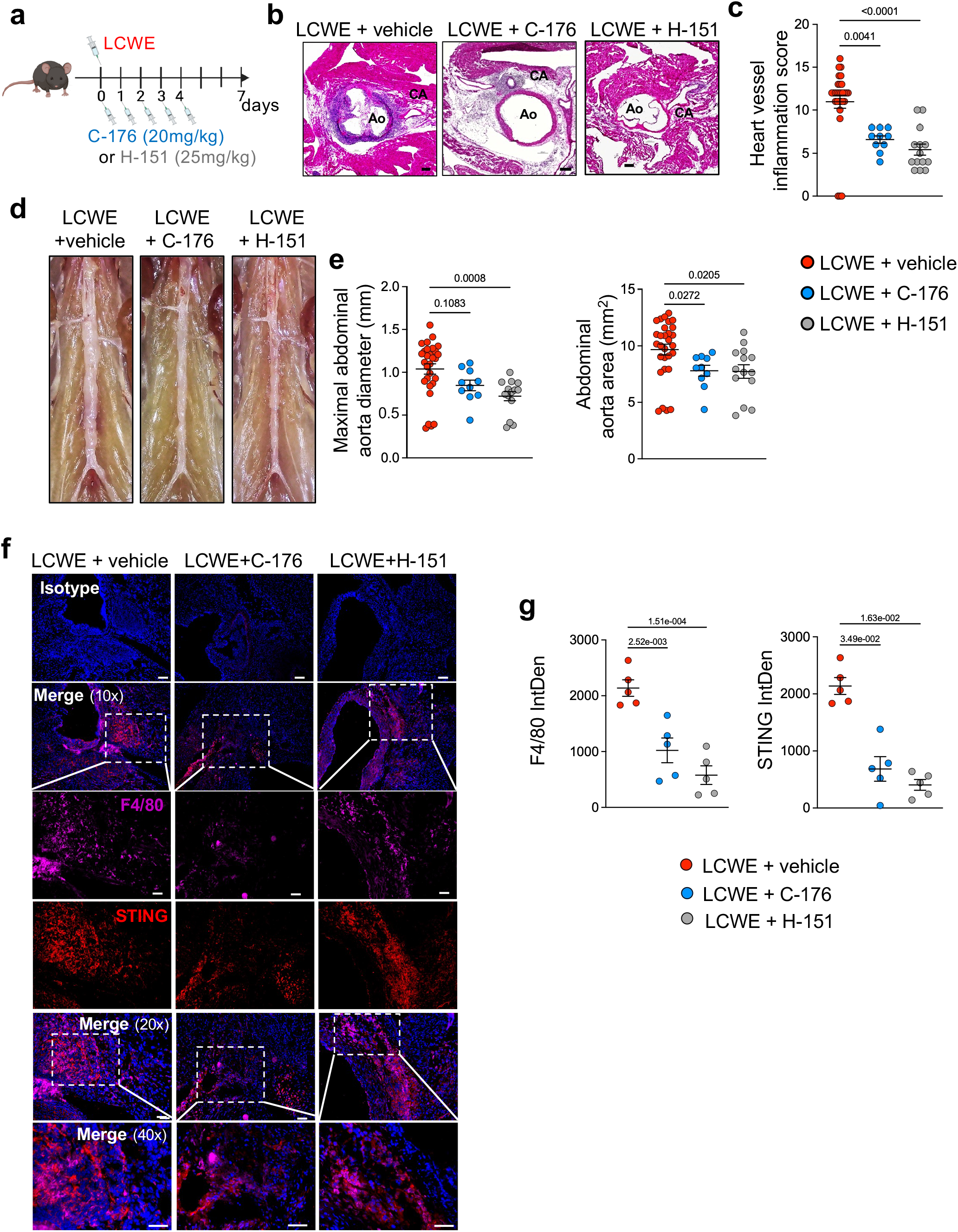
Pharmacological STING inhibition decreases the severity of LCWE-induced KD vasculitis. **a**, Schematic of the experimental design. WT mice were injected i.p. with either PBS or LCWE. Groups of LCWE-injected mice were treated with the STING inhibitors C-176 or H-151 daily from days 0 to 4 post-LCWE injection, and the severity of vasculitis was assessed at 1 week post-LCWE injection. **b, c,** H&E-stained heart sections (b) and heart vessel inflammation scores (c) of WT mice injected with either PBS, LCWE, LCWE and C-176, or LCWE and H-151, at 1 week post-injection (n = 10-25/group). Scale bars: 200 µm. **d, e,** Pictures of the abdominal aorta areas (d), maximal abdominal aorta diameter, and abdominal aorta area (e) measurements of WT mice injected with either PBS, LCWE, LCWE and C-176, and LCWE and H-151, at 2 weeks post-injection (n = 10-25/group). **f, g,** Immunofluorescent staining (IF) and integrated density (IntDen) quantification of STING and F4/80 (g) in heart tissues of WT mice injected with either LCWE, LCWE and C-176, or LCWE and H-151, at 2 weeks post-injection (n = 4-5/group). Data are presented as mean ± SEM. Pooled from 2 to 3 independent experiments (a-e). Each symbol represents one individual mouse. *p*-values obtained by Kruskal-Wallis with Dunn’s multiple comparisons test (c, e) and 1 way ANOVA with Tukey’s post-test (g).

## DISCUSSION

Although systemic immune responses associated with KD have been extensively investigated ^14–19,38^, the *in situ* immunopathogenic mechanisms leading to the development of CA aneurysms during KD are not well characterized. Here, we performed a multiplex spatial analysis of 64 proteins in rare CA tissue sections from KD and CTRL patients to determine the cellular landscape of affected CAs. We observed immune infiltrates in inflamed CAs and in the myocardium of KD patients. These infiltrations were characterized by a marked heterogeneity in both cellular composition and spatial distribution. Our neighborhood analysis indicates disrupted cellular organization across cardiac layers, expansion of immune neighborhoods, and contraction of structural compartments, with a pronounced reorganization of cellular and neighborhood interactions. While no significant enrichment of STING-related gene sets was detected in whole blood from patients with acute KD, we report a population of STING^+^ cells expressing CD45 and CD11b specifically in CA tissues from KD patients and identify the STING pathway as a previously unrecognized feature of KD pathogenesis. Using the LCWE-induced KD vasculitis murine model, which mimics key histopathological features of human KD ^8,9^, we observed increased expression and activation of the cGAS-STING pathway in inflamed CA tissues and abdominal aortic dilations from LCWE-injected mice. Furthermore, STING blockade, either genetic or pharmacological, reduced the development of LCWE-induced KD vasculitis, highlighting the cGAS-STING pathway as a promising therapeutic target to mitigate KD development and progression.

While KD mortality rate was initially estimated at 1 to 2% ^39^, it has significantly decreased (USA: 0.17% and Japan: 0.015% ^2^) with the introduction of IVIG and increased awareness of the disease among pediatricians. As a result, autopsies and collection of CA tissues from KD patients are extremely rare, and whole blood or PBMCs are more commonly used to characterize and phenotype KD systemic immune responses. Most transcriptomic, proteomic, and flow cytometric analyses of whole blood and serum from KD patients show immune dysregulation, enhanced innate immune responses, particularly neutrophils and monocytes, and elevated levels of IL-1β ^13,16,27,40,41^. scRNA-seq of PBMCs from patients with acute KD indicates upregulation of pro-inflammatory transcripts in monocytes and increased B cell abundance ^16^. These systemic alterations are amplified in patients who develop CA lesions, and such patients exhibit increased peripheral inflammatory cells, including monocytes, cytokine production, and complement activation ^19^. Although these studies have provided important insights into circulating immune responses during acute KD, the local immune mechanisms underlying and contributing to CA aneurysm formation remain poorly understood. A small number of prior studies have performed histopathological analyses of patient CA tissues and identified immune infiltrates composed predominantly of macrophages, neutrophils, CD8^+^ T cells, eosinophils, and IgA-producing plasma cells ^20,22–24,42–46^. Our multiplexed protein analysis of CA tissues from KD and CTRL patients provides an integrated, spatially resolved landscape of multiple cellular subsets at single-cell resolution *in situ* and not only confirms but also extends previous observations in their histological context. We validated the infiltration of macrophages, T cells, neutrophils, and DCs in the inflamed CA of KD patients. These infiltrates were localized near the inflamed CA vessel and persisted at greater distances from the lumen, extending into the myocardial tissue. Among key differences between CTRL and KD CA tissues, we detected IL-1β protein and STING^+^ cells, spatially localized to the inflamed epicardium and CA vessel wall. Our analysis indicates that KD is associated with increased presence of immune-rich neighborhoods, composed of macrophages and T cells, and our network analysis indicated enriched colocalization among immune cells, macrophages, T cells, and neutrophils in the epicardium and the CA vessel wall, whereas the bulk of enriched immune cell colocalizations in the myocardium were composed of neutrophils and CD4^+^ T cells. Beyond the expansion of these dense immune cell networks, we observed reduced colocalization between stromal cells (fibroblasts, VSMCs) and contraction of structural cellular compartments, including myocardial and endothelial layers, in CA tissues from KD patients.

Orenstein et al. proposed that three linked vasculopathic processes (acute necrotizing arteritis, subacute/chronic arteritis, and luminal myofibroblastic proliferation), which begin within the first 2 weeks of disease onset, may overlap rather than occur sequentially during KD ^7,22^. KD pathological processes involve neutrophils, lymphocytes, eosinophils, macrophages, and plasma cells infiltrating and progressively damaging the CA walls ^20,22,42,43,47^. Eventually, CA inflammation may progress to luminal myofibroblast proliferation, in which smooth muscle cell-derived myofibroblasts and their products progressively obstruct the CA lumen ^7,22^. In our cohort, macrophage and T cell densities were higher than neutrophil densities in the CA and myocardium. The KD CA tissues analyzed in this study were collected 18-33 days after disease onset, when necrotizing arteritis is expected to be resolved. Although neutrophils were detected in the CA and myocardium of KD patients, their low numbers may indicate overlap and heterogeneity among the three linked KD pathological processes. We also observed that immune cell infiltrations in the CA and myocardium of KD patients were associated with increased colocalization between immune cells. In contrast, colocalization of structural and stromal cells (VSMCs, nerve fascicles, fibroblasts, and myocardial cells) was reduced during acute KD. Together, our results indicate that KD coronary arteritis involves not only immune cells and tissue remodeling of the affected CA but also of the myocardium.

Among the CA tissues analyzed from patients with KD, the two tissue sections collected from different areas of the same KD patient (KD1a and KD1b) showed more intense infiltration of CD163^−^ macrophages in the CA and myocardium. In contrast, CA tissues from patients KD2 and KD3, both of which contained a thrombus in the affected CA, showed more extensive infiltration of CD163^+^ macrophages. Previous studies have reported CD163^+^ macrophages in CA tissues and aortic tissues from KD patients ^5,46^. CD163 is a scavenger receptor involved in the clearance of hemoglobin-haptoglobin complexes ^48^. In atherosclerosis, at sites of intraplaque hemorrhage, engulfment of these hemoglobin-haptoglobin complexes by CD163^+^ macrophages has been associated with production of proinflammatory cytokines (TNF-α, IL-1β, and IL-6), activation of ECs and VSMCs, and apoptotic programs that promote plaque instability ^49–51^. Our observation of higher densities of infiltrating CD163^+^ macrophages in KD tissue sections presenting with a CA thrombus may potentially reflect local hemolysis and formation of hemoglobin-haptoglobin complexes in these patients. The accumulation of hemoglobin-stimulated CD163^+^ macrophages and their production of proinflammatory cytokines could further exacerbate CA tissue damage. On the other hand, CCR2^−^ tissue-resident cardiac macrophages also express CD163, exhibit a less inflammatory phenotype, and are associated with resolution of inflammation and tissue repair programs ^52,53^. KD patients are treated with IVIG, which modulates immune responses and may influence the phenotype of tissue-infiltrating macrophages ^54^. Because our study did not assess hemolysis, measure hemoglobin-haptoglobin complexes, or allow deeper macrophage population phenotyping, further investigations are needed to determine the potential pro– or anti-inflammatory role of CD163^+^ tissue macrophages during KD.

Our analysis identified STING^+^ cells in CA tissues from KD patients. Furthermore, using the LCWE-murine KD model, we demonstrated activation of the cGAS-STING pathway in inflamed vascular tissues and showed that STING functionally contributes to LCWE-induced pathogenesis. STING is activated by cGAMP, which is produced by cGAS in response to cytosolic double-stranded DNA of microbial or endogenous origin ^55,56^. STING signaling induces type I IFN expression and activates NF-κB-dependent transcriptional programs ^55,56^. The NLRP3–IL-1β axis has been implicated in KD, as shown by increased expression of NLRP3– and IL-1β-related genes in whole blood ^13,18^, and higher circulating IL-1β ^28,29^ levels in patients with acute KD. Furthermore, several case reports indicate clinical improvement with anakinra, an IL-1 receptor antagonist, in IVIG-resistant KD patients ^57–61^. A functional crosstalk between the cGAS-STING and innate immune pathways, including the NLRP3 inflammasome, has been previously reported ^55^. Canonical NLRP3 inflammasome activation is a two-step process that requires priming followed by an activation signal ^62^. cGAMP, the second messenger of the cGAS-STING pathway ^63^, not only induces type I IFN signaling but also promotes inflammasome priming through the production of IFN-β ^64^. Furthermore, STING can recruit NLRP3 to the endoplasmic reticulum (ER) and activate it by attenuating its polyubiquitination in the context of HSV-1 infection ^65^. We detected both STING and IL-1β locally in CA tissues from patients with acute KD. However, their spatial overlap was limited, and STING and IL-1β mostly localized to adjacent, distinct tissue areas. While the cGAS-STING and NLRP3 inflammasome pathways may functionally converge ^55,62,65^, our analysis indicates that in inflamed CA from KD patients, their activation appears spatially compartmentalized. Host mitochondrial DNA (mtDNA), released into the cytosol during cellular stress and mitochondrial dysfunction, can activate both the cGAS-STING pathway and, in its oxidized form, the NLRP3 inflammasome ^66,67^. Altered mitochondrial function has been reported in both KD patients and the LCWE-induced KD model ^17,36,68–71^, often associated with reduced expression of mitochondrial and autophagy-related genes during the acute phase of the disease. Mitochondrial dysfunction may increase mtDNA accumulation and release, thus potentially activating both cGAS-STING and NLRP3 in parallel. While our previous studies indicated that impaired mitophagy and mitochondrial dysfunction contributed to LCWE-induced KD vasculitis in mice ^36,70^, it remains unclear whether STING activation in KD patients is driven by endogenous DNA released during mitochondrial stress and dysfunction, microbial DNA from an unidentified pathogen, or both.

Our study provides a spatially resolved, multiplexed analysis defining the landscape of inflamed CA tissues from patients with KD. Our analysis identifies multiple cell subsets and uncovers alterations in cellular composition and cell-to-cell localization within inflamed CA tissues from KD patients. We observed IL-1β deposition and the presence of STING^+^ myeloid cells, specific to inflamed CA tissues of KD patients. STING and IL-1β spatially localized in distinct tissue areas, and while STING blockade or genetic deficiency decreased the severity of LCWE-induced KD vasculitis, it did not completely prevent coronary arteritis and abdominal aortic dilations. Therefore, our results support further investigation on STING-targeted therapies, either alone or in combination with IL-1 blockade, as adjunctive therapies to IVIG.

## ACKNOWLEDGEMENTS

We thank Dr. Santiskulvong and the Cedars-Sinai Applied Genomics, Computation & Translational (AGCT) core for performing the 10x Visium HD spatial transcriptomics

## SOURCES OF FUNDING

M. Noval Rivas is supported by the NIH R01 HL139766. M. Arditi is supported by the NIH grants R01 HL170580 and NIH R21 AI193709. A.E. Atici is supported by the American Heart Association Career Development Award #1586278. This work was funded in part by Cedars-Sinai in support of CTSI grant UL1TR001881.

## AUTHOR CONTRIBUTIONS

M. Arditi, D. Zemmour, and M. Noval Rivas conceived the project and designed the experiments. B. Ross, A.E. Atici, Y. Lee, M.E. Lane, P. Ngai, and E. Schahrer performed experiments. K. Takahashi, Y. Yokouchi, and M.F. Fishbein provided pathological samples and pathological review. B. Ross, P. Ngai, and D. Zemmour designed the bioinformatic pipeline and performed the bioinformatics analysis. B. Ross, A. E. Atici, Y. Lee, D. Zemmour, and M. Noval Rivas analyzed the data. B. Ross, M. Arditi, K. Takahashi, D. Zemmour and M. Noval Rivas interpreted the results. B. Ross, D. Zemmour, and M. Noval Rivas wrote the manuscript with input from all authors.

## DISCLOSURES

The authors report no conflict of interest.

## DATA AND CODE AVAILABILITY

Spatial omics data used in this study are available at the Gene Expression Omnibus database under GSE350998 and GSE350866. The code used for the bioinformatic analyses is available on GitHub (https://github.com/NovalRivasLab/Kawasaki-Disease-Spatial).

## MATERIALS AND METHODS

### Patient cohort

Previously collected, archived, and de-identified formalin-fixed, paraffin-embedded (FFPE) human heart tissue sections and limited associated clinical data were obtained from Toho University, Japan, and the University of California, Los Angeles (UCLA). Tissues were collected under protocols approved by the Ethics Committee of Toho University Ohashi Medical Center (Approval no. 00003131) and the UCLA Institutional Review Board (protocol no. IRB11-002516-AM-00007). KD cases were diagnosed according to the Japanese guidelines in effect at that time. All specimens were provided de-identified. Cedars-Sinai Medical Center Institutional Review Board Office determined that this study did not constitute human subjects research. Coronary artery (CA) tissues were obtained from 3 patients who died from KD, yielding a total of 4 analyzed tissue sections, including 2 specimens prepared from different tissue blocks of the same patient (KD1a and KD1b). Control (CTRL) tissues were obtained from 2 different patients who died from either a congenital heart disease that did not involve the CA or meningitis, yielding a total of 3 analyzed CTRL tissue sections, including 2 sections from the same patient (CTRL2a and CTRL2b). **Supplementary Table 1** summarizes patient demographics, disease duration, treatment history, and pathological findings. Hematoxylin and eosin (H&E) staining was used for routine histological and pathological examination, and elastic Verhoeff-van Gieson (EVG) staining was performed to visualize elastic tissue structures.

### High-resolution spatial proteomic analysis of human heart tissues

#### CosMx SMI Protein Assay

The CosMx Spatial Molecular Imager (SMI) (Bruker Spatial Biology, Bothell, WA) ^72^ was used for spatial protein profiling of FFPE heart tissue sections (n = 7 samples; total slides = 6). FFPE tissue sections were cut at 5 µm thickness and mounted on 6 slides. Following the manufacturer’s instructions (CosMx SMI Manual Slide Preparation for Protein Assays, MAN-10185-03; CosMx SMI Instrument User Manual, MAN-10161-08), slides were processed using the CosMx SMI Protein FFPE assay. Briefly, sections underwent routine FFPE immunostaining preparation, including baking, deparaffinization, rehydration, antigen retrieval in 1X CosMx Target Retrieval Solution, and blocking in Buffer W. For high-plex spatial protein detection, slides were incubated overnight at 4°C with the 64-plex CosMx protein antibody panel, together with CD298/B2M segmentation marker, PanCK/CD45 supplemental marker, and CD3 à la carte marker. Slides were then washed and processed with CosMx fiducials for image registration, post-fixed with 4% paraformaldehyde, stained with DAPI nuclear stain, and treated with NHS-acetate according to the manufacturer’s protocol. Flow cells were assembled and loaded onto the CosMx SMI instrument for cyclic reporter binding and fluorescence imaging. Run setup used pre-bleaching profile C, with DAPI for nuclear segmentation, B2M/CD45/PanCK/CD3 for membrane segmentation, and the Immuno-Oncology 64-plex protein core panel. The results from these runs were then uploaded to the AtoMx Spatial Informatics Platform (Bruker Spatial Biology). Data for all 6 CosMx SMI runs, including protein count matrices, FOV positions, cell segmentation coordinates, and metadata, from AtoMx for further analysis.

#### Tissue Imaging

Protein fluorescence and cell segmentation images were produced in napari ^73^ and edited in GNU Image Manipulation Program (GIMP).

#### Filtration, Normalization, and Dimensional Reduction

Data for all 6 CosMx SMI runs, including protein count matrices, FOV positions, cell segmentation coordinates, and metadata, were downloaded from Bruker’s AtoMx Spatial Informatics Platform (Bruker Spatial Biology). These were then loaded into R (v4.5.1) via package Seurat ^74^ (v5.3.0) to produce Seurat objects for further downstream analysis. For each slide, distorted tissue regions, such as CA thrombi (KD2, KD3), folded, torn, and detached tissue, and regions of protein hyperfluorescence/artifacts were removed first. Segmented cells outside of the tissue region, with abnormally large surface area (>500 µm^2^), high background protein expression (i.e., high Rabbit (Rb) IgG and/or Mouse (Ms) IgG1 fluorescence intensity), and/or high total protein expression (>20,000 combined protein fluorescence intensity) were then removed from the slides. The slides’ protein assays were normalized via centered log ratio (CLR) normalization within individual FOVs via Seurat’s NormalizeData function (normalization.method = “CLR”, margin = 1). Variable features were selected based on protein fluorescence quality and assigned with the VariableFeatures function, and the normalized data were then scaled using ScaleData. Selected proteins for each slide are described in **Supplementary Table 2.** Dimensionality reduction was then performed with RunPCA. The first 30 principal components (except for KD3, in which 27 were used as only 27 proteins were selected as variable features) were used to construct the Uniform Manifold Approximation and Projection (UMAP) for each slide with the RunUMAP function. These principal components were also used for neighbor graph construction via FindNeighbors, which was utilized for unsupervised Louvain clustering of the cells, calculated by FindClusters (resolution = 4, method = “igraph”).

#### Tissue Region and Cell Type Annotation

Tissue region annotations were assigned to cells, utilizing the H&E and napari-generated imaging for reference. To identify cell populations in the samples, differentially expressed proteins (log_2_-Fold Change (FC) >0.5 and adjusted *p*-value <0.05) were calculated for the clusters via FindMarkers (test.use = “wilcox”, only.pos = T). Cells were first annotated for putative cell types based on the clusters’ differential expression of proteins. Cells in the epicardium and vessel wall in unannotated clusters were annotated via FindTransferAnchors (reduction = ‘pcaproject’) and TransferData, utilizing cells from annotated clusters in the epicardium and vessel wall as the reference data to project annotations onto the unannotated cells. Cells in the myocardium from unannotated clusters were labeled myocardial cells. Across all 6 samples, 20 major cell populations were identified, including stromal cells (ECs, VSMCs, EC/VSMCs, fibroblasts), myeloid cells (CD163^+/–^ macrophages, dendritic cells, neutrophils), and lymphocytes (B cells, naïve/activated/regulatory CD4^+^ T cells, naïve/activated CD8^+^ T cells, other T cells, and natural killer (NK) cells), as well as nerve cells, myocardial cells, CD38^+^/CD39^+^ cells, and STING^+^ cells. Spatial images for the slides were plotted with ImageDimPlot.

#### Slide Integration

Slides were concatenated into a singular Seurat object, and principal components were recalculated via RunPCA for the combined object based on all proteins except background features Rb IgG and Ms IgG1. The slides were then integrated via the R package Harmony ^25^ based on the concatenated object’s principal components with the cells’ sample ID and FOV number as metadata variables. An integrated UMAP for the slides was calculated based on the first 30 principal components calculated by the Harmony integration ^25^. The integrated UMAP was plotted with DimPlot, and cell type markers were recalculated for validation in the integrated object with FindMarkers (test.use = “wilcox”). Expression of key markers differentiating cell types were plotted using ggplot2 ^75^.

#### Cell Type Statistical Analysis

For statistical analysis of cell type populations in KD vs. control samples, two serial sections from CTRL2 were separated into CTRL2a and CTRL2b, totaling 7 sections analyzed (4 KD, 3 control). For statistical comparison of the different regions of the heart in KD vs. control, samples were separated into the epicardium/vessel wall and myocardium regions based on H&E staining and CosMx imaging topography. Cell type proportions in sample regions were calculated by dividing the cell type’s population by the entire region’s cell count. To calculate cell type density, the area of each sample’s regions was estimated with alphahull ^76^, and the cell type’s population was divided by the calculated area. Cell type densities and their corresponding proportions in each sample were visualized by region in a heatmap created with the R package pheatmap ^77^. Cell type densities were also visualized in bar plots comparing KD vs. control samples with ggplot2 ^75^. Significance of cell type densities in the KD vs. the control samples was calculated on the log_10_-transformed densities via the R package stats t.test function (alternative = ‘greater’, var.equal = F, paired = F). Instantaneous cell type density at the cell level was calculated by summing the number of neighboring cells of the same type in the 1 mm^2^ area surrounding the centroid of a given cell. These instantaneous densities for cells of interest were plotted on the entire sample image and log_10_-transformed for visualization in ridge plots via ggplot2^75^.

#### Cell Pair Enrichment

In the epicardium/vessel wall and myocardium of the KD and control samples, the closest neighboring cells within 20 µm for each cell were identified via R package RANN’s nn2 function (k=2, radius = 20) ^78^ and summed by cell type pairs. For analysis of enriched cell type pairs in KD vs control samples, cell pair counts were normalized by trimmed mean of M-values (TMM) normalization and then log-transformed by counts per million (CPM) with a prior count of 0.01. Differential enrichment was then calculated by fitting a linear model matrix to the normalized KD vs. control cell type pair counts via limma-trend’s “lmFit” method ^79^, and adjusted with empirical Bayes statistics via edgeR’s eBayes function ^80^. Differentially enriched cell type pairs in KD vs. control samples were identified by a Benjamini-Hochberg False Discovery Rate (FDR) <0.05, which were visualized in a circle plot with the R package fields ^81^.

#### Neighborhood Analysis

Samples were projected onto a singular spatial plane within the Harmony-integrated ^25^ Seurat object to identify cellular neighborhoods across all KD and control samples. Seurat’s BuildNicheAssay (neighbors.k = 5, niches.k = 9) ^74^ was then run to calculate 9 distinct neighborhoods. The distribution of niches was visualized in each sample via ImageDimPlot. Proportions of cell types present in each neighborhood were calculated by dividing the cell type population by the total count of cells present in the neighborhood. For proportions of samples present in each neighborhood, the number of a sample’s cells present in a neighborhood were divided by the total count of cells present in the neighborhood, calculated by region (epicardium/vessel wall and myocardium). Proportions for both cell types and samples in the niches were visualized via pheatmap ^77^. The distribution of neighborhoods was separated by observation of contraction or expansion in the KD vs. control samples and individually by neighborhood to be visualized in each sample via ImageDimPlot. Following the same analysis for cell pair enrichment, neighboring cells within 20 µm for each cell were identified via R package RANN’s nn2 function (k = 2, radius = 20) ^78^ and summed by neighborhoods. For analysis of enriched cell neighborhood pairs in KD vs control samples, cell neighborhood pair counts were normalized by trimmed mean of M-values (TMM) normalization and then log-transformed by counts per million (CPM) with a prior count of 0.01. Differential enrichment was then calculated by fitting a linear model matrix to the normalized KD vs. control cell neighborhood pair counts via limma-trend’s “lmFit” method ^79^, and adjusted with empirical Bayes statistics via edgeR’s eBayes function ^80^. Differentially enriched cell neighborhood pairs in KD vs. control samples were identified by a Benjamini-Hochberg FDR <0.05, which were visualized in a circle plot with the R package fields ^81^. Density of neighborhoods in the epicardium/vessel wall and myocardium for each sample was calculated by dividing the population of cells in the given neighborhood by the alphahull-estimated area of the region ^76^. To determine differential neighborhood abundance by region between KD and control samples, the fold change of the average neighborhood density of KD samples relative to the average neighborhood density of control samples was calculated for each neighborhood and then log_2_-transformed for both the epicardium/vessel wall and myocardium regions. A Shapiro-Wilk test was performed on the KD and control samples separately to assess the normality of the density distributions via the R package stats’ shapiro.test function. The function t.test (alternative = “two-sided”, var.equal = T) was used for normally distributed neighborhood densities (Shapiro-Wilk *p*-value >0.05) to assess the significance of the neighborhood’s differential abundance, and a Wilcoxon Rank Sum test was performed using the R package stats’ wilcox.test function (alternative = “two-sided”) when neighborhood densities significantly deviated from normality (Shapiro-Wilk *p*-value <0.05). Results of the differential abundance calculations were visualized in a dot plot via ggplot2 ^75^.

#### IL-1β Cell Population Analysis

Cells with a CLR-normalized expression of IL-1β >1.5 were identified as phenotypically IL-1β^+^ cells within their annotated cell type. The instantaneous density of these cells was calculated by summing the number of neighboring IL-1β^+^ cells in the 1 mm^2^ area surrounding a given cell. These densities were plotted on the entire sample image as well as the proportion of IL-1β^+^ cells by cell type via ggplot2 ^75^.

#### STING^+^ Cell Population Analysis

STING^+^ cells were isolated from the concatenated Seurat object for downstream analysis. Principal components were recalculated via RunPCA for the combined object, excluding rabbit IgG and Ms IgG1. The cells were integrated with Harmony ^25^ with the same specifications as the concatenated slides, with the corresponding UMAP recalculated accordingly as well. Expression of proteins were visualized utilizing the scaled protein data on the new UMAP via DimPlot.

#### Lumen Distance

To quantify immune cell infiltration in the tissue surrounding the CA lumen in the KD samples, cells bordering the CA lumen were selected, and their coordinates in mm were extracted and converted into a simple feature object with the R package sf ^82^. All cells and corresponding coordinates (in mm) were converted into a simple feature object, and the minimum Euclidean distance between each cell and the cells bordering the lumen were calculated the via sf’s st_distance function ^82^. The minimum distance from any lumen was extracted, with cells with a minimum distance ≥ 2 mm filtered out. Cells ≤ 2 mm from the lumen were visualized spatially on the samples as well as the proportion by cell type and neighborhood via ggplot2 ^75^.

### Gene set enrichment analysis of publicly available human whole blood and murine abdominal aorta gene expression datasets

A curated gene panel combining genes related to cGAS-STING signaling (KEGG Cytosolic DNA-sensing pathway; Ishikawa STING signaling; Reactome STING mediated induction of host immune responses; GOBP cGAS/STING signaling pathway; GOBP negative regulation of cGAS/STING signaling pathway; BioCarta NF-Kb Signaling Pathway), inflammatory responses (Reactome Interleukin-1 family signaling; Reactome The NLRP3 inflammasome; GOCC NLRP3 inflammasome complex; KEGG MEDICUS Reference NLRP3 inflammasome signaling pathway), and type I interferon response (GOBP response to type I interferon; GOBP cellular response to interferon alpha; GOBP response to interferon alpha; Hallmark interferon alpha response; GOBP cellular response to interferon beta; GOBP response to interferon beta; ST Type I Interferon Pathway) were accessed from MSigDB ^83^ Human Collections H (hallmark gene sets), C2 (curated gene sets), and C5 (ontology gene sets) (**Supplementary Table 3**). Altogether, these pathway gene sets contained 392 non-redundant genes. We then analyzed expression of these genes in publicly available whole-blood gene expression datasets from healthy controls (HC), febrile controls (FC), patients with acute KD, and IVIG-treated convalescent KD patients. Whole blood microarray dataset GSE68004 ^15^was accessed from Gene Expression Omnibus (GEO) and normalized and analyzed for expression of genes in acute KD vs. HC samples via the R package limma ^79^. Whole blood bulk RNA dataset GSE178491 ^30^ was also accessed via GEO and analyzed for gene expression in acute KD vs. convalescent KD and FC samples using the R package DESeq2 ^84^. The gene expression dataset generated from the abdominal aorta of PBS– and LCWE-injected mice GSE141072 ^34^ was also accessed via GEO and analyzed for gene expression using the R package DESeq2 ^84^. Gene expression in these datasets was then analyzed for enrichment of all available gene sets in MSigDB ^83^ Human Collections H, C2, and C5 via R package fgsea’s ^85^ fgseaMultilevel function. Enrichment of pathways in the curated panel was visualized with the plotGseaTable function for each dataset.

### Mice

WT C57BL/6J mice were obtained from the Jackson Laboratory. *Sting^gt/gt^* mice (Goldenticket or *Tmem173^Gt^*, Jackson Laboratory Strain #017537) were a kind gift from Dr. Sutterwala at Cedars-Sinai Medical Center. Only male animals were used in this study, as LCWE injection induces stronger and more consistent coronary vasculitis and abdominal aorta dilation in male mice than in female mice ^8,34^. Mice were randomly assigned to groups, housed under specific pathogen-free conditions, and used in accordance with the Cedars-Sinai Medical Center Institutional Committee guidelines (IACUC009436).

### The Lactobacillus casei cell wall extract (LCWE)-induced murine model of KD vasculitis

LCWE was prepared as previously published ^9^. Five-week-old male mice were injected intraperitoneally (i.p.) with 500 µL of LCWE or PBS (control group). After one to two weeks, as indicated in the figure legends, mice were euthanized, perfused with PBS, and hearts and abdominal aortas were collected. After dissection, abdominal aorta tissues were photographed to calculate maximal abdominal aorta diameter by measuring 5 areas separated by 2 mm along the infrarenal abdominal aorta (below the left renal artery) and measuring the infrarenal abdominal aorta area (ImageJ; NIH)^8,13^. Heart and abdominal aorta tissues were embedded in O.C.T., and serial cryosections (7 µm) were prepared for hematoxylin and eosin (H&E, Sigma-Aldrich) staining and immunofluorescence (IF). Histopathological examination and inflammation severity scoring of heart tissues were performed on H&E-stained sections. Heart vessel inflammatory score (coronary arteritis, aortic root vasculitis, and myocarditis) was determined using a previously published score ^9,86^. Only heart sections showing an inflamed CA branch separating from the aorta were scored. Briefly, acute inflammation, chronic inflammation, and connective tissue proliferation were each assessed using the following scoring system: 0 = no inflammation; 1 = rare inflammatory cells; 2 = scattered inflammatory cells; 3 = diffuse inflammatory cell infiltrate; and 4 = dense clusters of inflammatory cells. Fibrosis was determined using the following scoring system: 0 = no medial fibrosis, 1 = medial fibrosis involving less than 10% of the CA circumference, 2 = medial fibrosis involving 11% to 50% of the CA circumference, 3 = medial fibrosis involving 51% to 75% of the CA circumference, and 4 = medial fibrosis involving more than 75% of the CA circumference. All 4 scores were combined to generate a severity score called “Heart inflammation score”, as published ^9,86^. Pictures were taken using a Biorevo BZ-9000 or BZ-X710 microscope (Keyence).

### Single-cell RNA sequencing (scRNA-seq) of murine abdominal aortas

The murine scRNA-seq dataset GSE178765 ^13^, generated from abdominal aortas of PBS– and LCWE-injected mice, was accessed via the GEO database. Counts were normalized with NormalizeData (normalization.method = ‘LogNormalize’) and then scaled from 0 to 1. The expression of *Sting1* was visualized on the calculated UMAP via DimPlot.

### Spatial gene expression of heart tissues from PBS– and LCWE-injected mice (Visium HD)

RNA was extracted from fresh-frozen heart tissue scrolls of PBS– and LCWE-injected mice (n = 5/group) using the *Quick*-RNA Miniprep kit (Zymo Research, Irvine, CA) and analyzed for RNA integrity on the Agilent 4200 TapeStation (Agilent Technologies, Santa Clara, CA). All samples had an RNA Integrity Number (RIN) of 4 or higher. Fresh-frozen heart tissues were sectioned at 10 µm, adhered to pre-chilled glass slides, frozen, and placed in a pre-chilled slide mailer on dry ice, as per the Visium HD Fresh Frozen Tissue Preparation Handbook (10x Genomics CG000763). Slides were stored at –80 °C. To assess tissue quality, sections were fixed in 3.7% formaldehyde and stained with DAPI to examine nuclear morphology and with H&E to assess tissue morphology and block quality. Based on RIN measurements and H&E-based histological assessment, one qualified tissue section from each group was prepared for the Visium HD assay in accordance with the Visium HD Fresh Frozen Tissue Preparation Handbook. Tissue sections on glass slides were fixed with formaldehyde, stained with H&E, and cover-slipped with glycerol and imaged on an Aperio AT2 (Leica Biosystems, Buffalo Grove, IL) at 20X and 40X magnification. The coverslip was then removed from the slides, and tissues were destained and permeabilized, then hybridized with Mouse WT Probes v2 for 17.5 hours according to the Visium HD Spatial Gene Expression Reagent Kits User Guide (10x Genomics CG000685). Post-hybridization washes were followed by probe ligation and wash, and the ligated probes were released and transferred onto Visium spatially barcoded slides using the Visium CytAssist instrument (10x Genomics, Pleasanton, CA). Probe extension was performed on the ligated product, followed by pre-amplification and cleanup. A portion of the pre-amplification product was used in a qPCR reaction to determine the sample index PCR cycle number. A final PCR amplification and cleanup generated indexed libraries. Multiplexed libraries were quantified via the Qubit 1X dsDNA HS Assay kit (ThermoFisher Scientific, Waltham, MA), and library size was determined on the 4200 TapeStation (Agilent Technologies) with the HS D1000 Screen Tape. Libraries were sequenced on a NovaSeq X Plus (Illumina, San Diego, CA) with sequencing configuration 43×10×10×50 bp and a sequencing depth of 550,000,000 reads/6.5×6.5 mm capture area.

#### Data Analysis

Visium HD transcriptomic profiling was performed on one heart tissue section from a PBS-injected control mouse, and one heart tissue section from an LCWE-injected mouse. The sections were analyzed with Space Ranger count (v4.0) (10X Genomics), where the samples’ FASTQ files were mapped to the murine gene reference probe set GRCm39 2024-A, and transcripts were mapped to the CytAssist and 40X microscope image. Binned outputs at the 16 µm resolution were utilized for downstream analysis of gene expression and were converted into AnnData objects with the Python library SpatialData ^87^. Distorted regions, such as folded/torn tissue, detached tissue, and bins not mapped to tissue, were removed from both samples. Bins were then log-normalized via Python library SCANPY’s normalize.total (target_sum = 1e6, copy = True) and log1p functions ^88^. For analysis of *Sting1* (STING), *Ptprc* (CD45), and *Itgam* (CD11b) expression, the two samples were concatenated into a singular AnnData object based on overlapping genes. Normalized transcripts were then scaled 0 to 1 across the samples, and selected genes were visualized in the LCWE samples vs. the PBS samples with Python library Squidpy’s spatial_scatter plotting function ^89^. To visualize *Sting1* and *Il1b* codetection, bins were labeled as *Sting1^-^Il1b^-^, Sting1^+^Il1b^-^, Sting1^-^Il1b^+^,* or *Sting^+^Il1b^+^*based on absence or presence of normalized transcripts counts in the given samples’ bins, which were visualized in the LCWE sample vs. the PBS sample with Squidpy’s spatial_scatter plotting function ^89^.

### Protein lysates, SDS/PAGE electrophoresis, transfer, and Western blotting of heart tissues and abdominal aortas from PBS– and LCWE-injected mice

Heart and abdominal aorta tissues from PBS– and LCWE-injected mice were lysed in RIPA buffer (10X stock: 0.5 M Tris-HCl, pH 7.4, 1.5 M NaCl, 2.5% deoxycholic acid, 10% NP-40, 10 mM EDTA, Millipore; 20-188) with 1X phosphatase inhibitor cocktail 3 and 1X protease inhibitor cocktail (Sigma-Aldrich; P0044 and P8340). Lysates were cleared by brief centrifugation (10 min at 8000g), normalized, and boiled at 95 °C after the addition of 6X SDS protein loading dye. Proteins were then loaded onto a gradient SDS/PAGE gel and transferred to nitrocellulose or PVDF membranes. Membranes were blocked (1 hour at room temperature) and incubated overnight at 4°C with primary antibodies from the Mouse-Reactive STING pathway Antibody Sampler kit (Cell Signaling Technology, #16029T), followed by incubation with appropriate secondary antibodies (1 hour at room temperature) in 5% (w/v) dry milk or bovine serum albumin (in tris-buffered saline buffer with 0.1% Tween-20 (v/v)) on a shaker. Membranes were developed in ECL prime reagent (ThermoFisher Scientific; 34577), and images were captured with ChemiDoc Imager (BioRad). Quantifications were performed with Image Lab software (Bio-Rad). Protein expression was quantified by densitometry and normalized to either Ponceau S. or β-actin. For samples with no detectable band after background subtraction, a “0” value was assigned.

### Immunofluorescence staining

OCT frozen mouse heart tissue serial sections (7 µm) were fixed in ice-cold acetone for 5 minutes, washed with PBS-Tween (PBS-t; 0.05% Tween-20), and blocked for 1 hour at room temperature with serum-free protein block (Agilent; #X090930-2). Sections were then incubated with primary antibodies (STING; Proteintech #19851-1-AP or F4/80; Cytek Biosciences #704801U100) or isotype controls in a 20% serum-free protein block in PBS-t overnight at 4°C in a humid chamber. The next day, sections were washed with PBS-t, incubated 1 hour at room temperature with fluorescently labeled secondary antibodies (anti-rabbit Alexa Fluor 647 or anti-rat Alexa Fluor 555) 20% serum-free protein block in PBS-t. After incubation, sections were washed again and mounted with Fluoroshield mounting medium with DAPI (Abcam; ab104139). Images were obtained using a Biorevo BZ-X710 (Keyence) fluorescent microscope and were analyzed with ImageJ (NIH) software. For all immunofluorescent staining, we used isotype controls for primary antibodies to distinguish genuine target staining from background.

### IL-1β quantification from mice peritoneal lavage

Mice were injected with either PBS or LCWE as described above. 24 hours later, we collected peritoneal lavage in sterile PBS, centrifuged at 400 g for 5 minutes at 4 °C, and discarded the cells. IL-1β concentration was measured in the supernatants by ELISA (R&D Systems, Mouse IL-1 beta/IL-1F2 DuoSet ELISA; DY401) according to the manufacturer’s protocol ^36^.

### STING inhibition during LCWE-induced KD

WT mice were injected i.p. with vehicle (5% DMSO, 5% PEG300, 5% Tween 80, 85% DW), C-176 (20 mg/kg of body weight; Catalog. # T5154; TargetMol Chemicals Inc.), or H-151 (25 mg/kg; of body weight; Catalog. #T5674; TargetMol Chemicals Inc.) diluted in vehicle. Mice received the STING inhibitors daily from days 0 to 4 post-LCWE injection. The severity of vasculitis was assessed 1 week post-LCWE injection, as described above.

### Statistical analysis and software

GraphPad Prism 10 or 11 (GraphPad Software, Inc.) and R software were used for statistical analysis and graph creation. No statistical methods were used to predetermine the sample size. The statistical test used for each comparison is indicated in the corresponding figure legends. Normality of the data was determined with a Shapiro-Wilk test (α = 0.05). For comparisons of 2 groups, one or two-tailed unpaired t-tests were used for normally distributed data, with Welch’s correction for unequal variances when necessary. For nonparametric data, the Mann-Whitney test was used. For more than 2 group comparisons, one-way ANOVA with Tukey’s post hoc test was used for normally distributed data. The Kruskal-Wallis test with Dunn’s multiple comparisons test was used for non-normally distributed data. Results are reported as means ± SEM, where each point represents one sample. A *p*-value <0.05 was considered statistically significant. Schematics used in the figures were created with BioRender.com. ChatGPT (OpenAI) was used for language editing and code optimization.

---

**Supplementary Table 1.** Characteristics of coronary artery (CA) tissue from patients with Kawasaki disease (KD) and controls (CTRL).

| Kawasaki disease (KD) |  |  |  |  |  |  |
| --- | --- | --- | --- | --- | --- | --- |
| ID | Age (months) | Sex | Illness duration (days) | Treatment | Pathology | Cause of death |
| KD1a<br>KD1b | 5 | Male | 33 | IVIG, Aspirin | Coronary arteritis with aneurysms in the LCA (KD1a) and RCA (KD1b).<br><u>KD1a</u>: LCx with intact internal elastic lamina. Transmural cellular infiltration, predominantly composed of macrophages and a few lymphocytes, as well as mild pericardial inflammation.<br><u>KD1b</u>: Dilation with circumferential fibrocellular intimal thickening of the RCA. Mild interstitial and perivascular infiltrations were also observed in the myocardium and pericardium adjacent to the RCA. RCA medial smooth muscle cells were lost, and both the internal and external elastic laminae are fragmented. Macrophage infiltration was observed throughout the full thickness of the vessel wall. | KD |
| KD2 | 11 | Male | 18 | IVIG, Aspirin | Coronary arteritis with a giant aneurysm (8 mm diameter) in the RCA with a fresh thrombus in the RCA lumen.<br>Marked media thinning, disruption of both the internal and external elastic laminae, and transmural inflammatory infiltration predominantly composed of macrophages with a few lymphocytes. | KD |
| KD3 | 3 | Female | 23 | Unknown | Coronary arteritis with an aneurysm of unknown location, potentially in the LCA, and an organized thrombus.<br>Disruption of the CA wall was observed, with marked media thinning and absence of the internal and external elastic laminae.<br>A macrophage-predominant transmural inflammatory infiltrate with a few lymphocytes was noted, as well as mild infiltration of immune cells in the myocardium. | KD |
| Control (CTRL) |  |  |  |  |  |  |
| ID | Age (months) | Sex | Illness duration (days) | Treatment | Pathology | Cause of death |
| CTRL1 | 6 | Male | -- | -- | Normal CA | Complex congenital heart disease |
| CTRL2a<br>CTRL2b | 12 | Female | -- | -- | Normal CA | Meningitis |

Abbreviations: CA, coronary artery; CTRL, control; IVIG, intravenous immunoglobulin; KD, Kawasaki disease; LCA, left coronary artery; LCx, left circumflex; RCA, right coronary artery.

**Supplementary Table 2.**
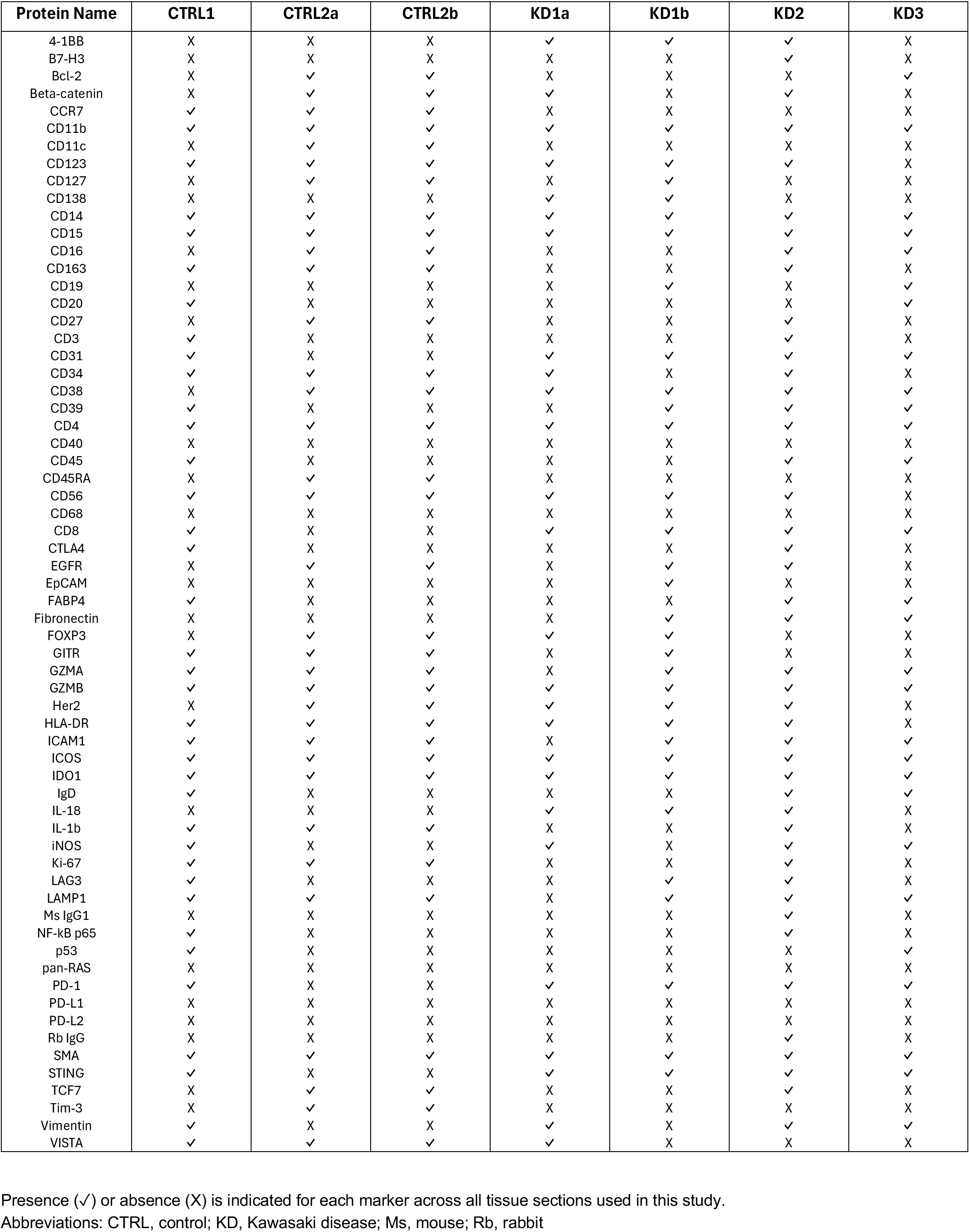
Marker expression across coronary artery tissue sections.

**Supplementary Table 3.**
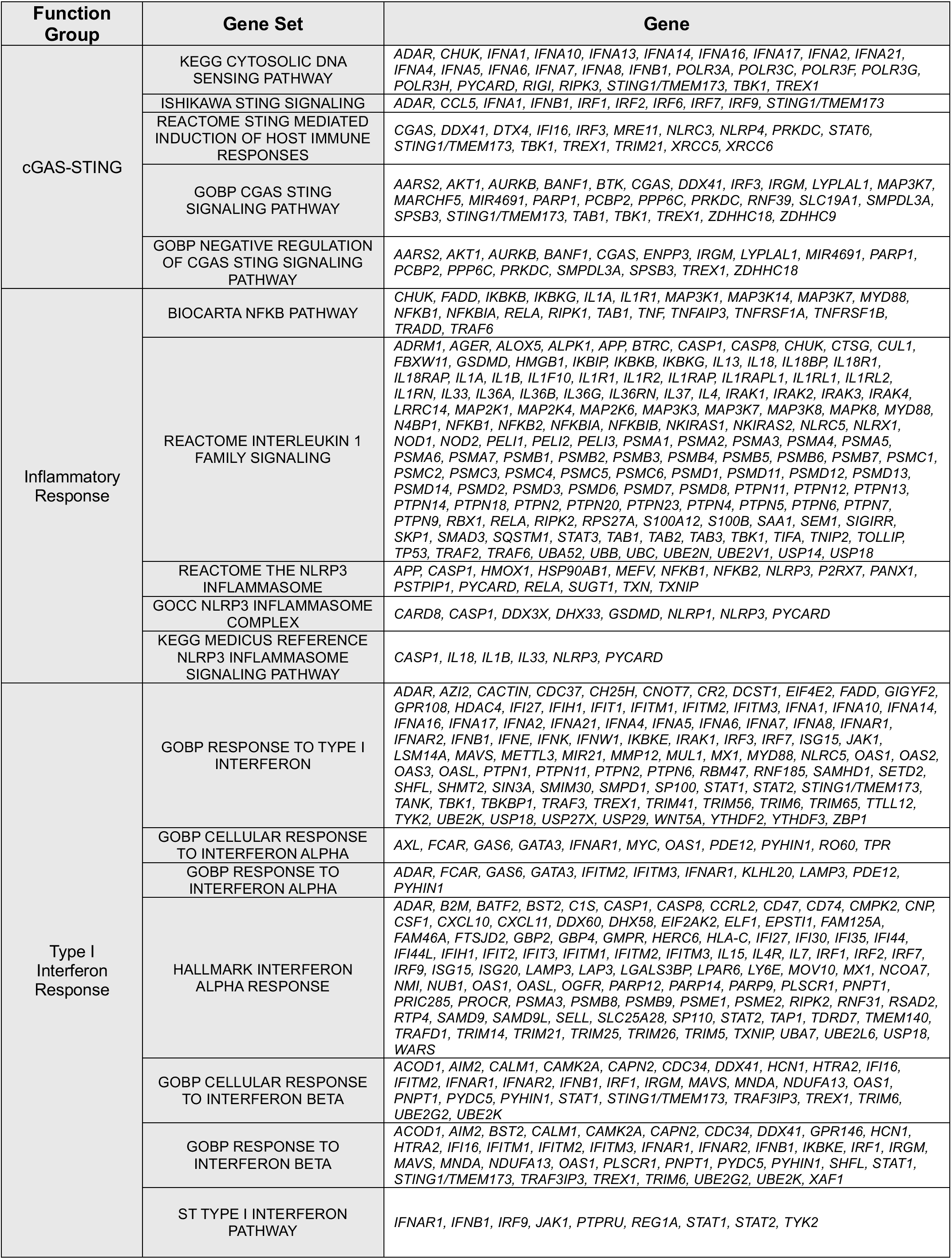
– List of selected pathways and genes used for gene expression analysis.

## Notes

### Competing Interest Statement

The authors have declared no competing interest.

