## Supplementary Figures for "Spatial profiling of Kawasaki disease cardiac inflammation identifies STING as a therapeutic target"

### Extended Data Figure 1

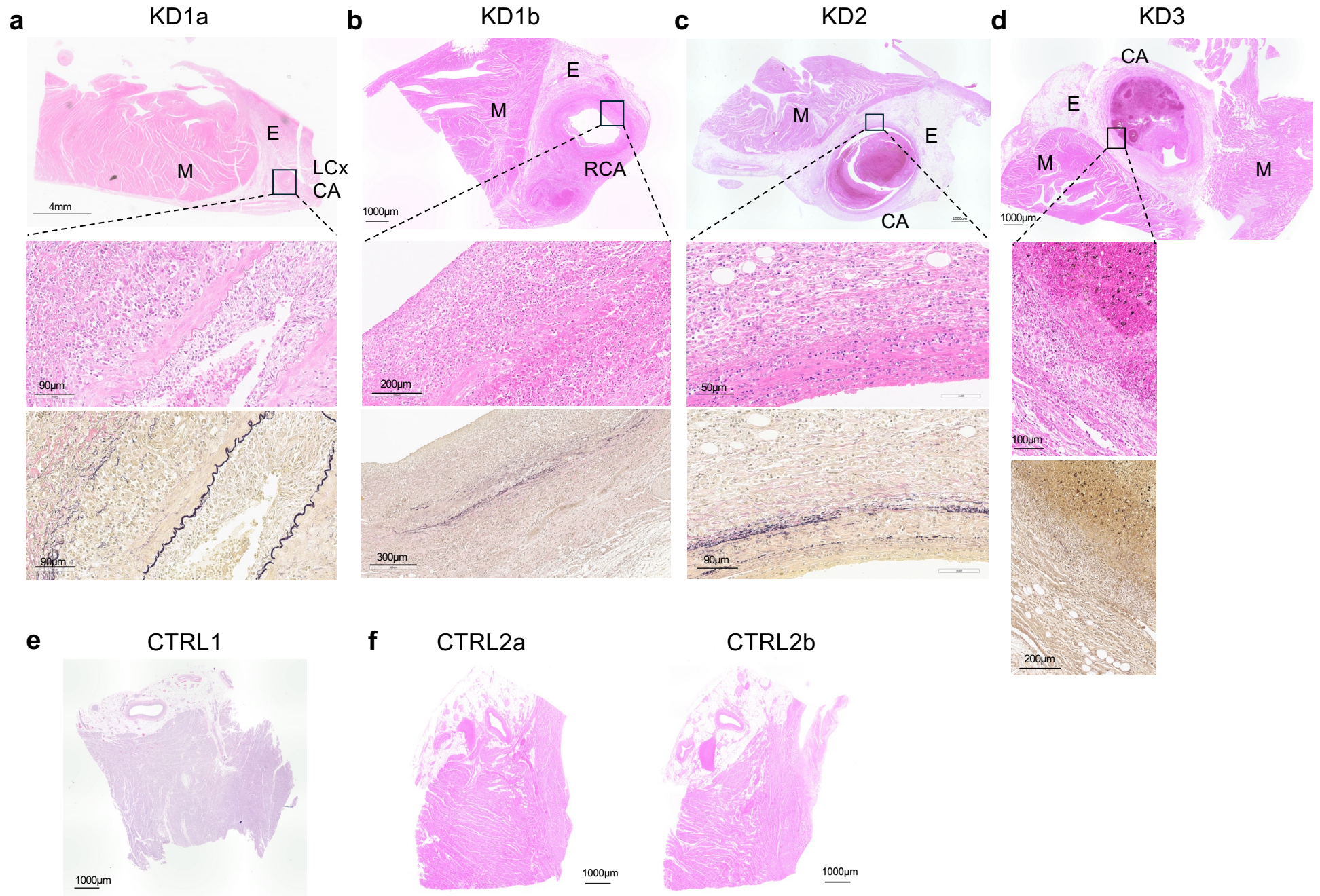

**Extended Data Figure 1. Histological analysis of coronary artery tissues from control and Kawasaki disease patients. a–d,** Microscope images showing hematoxylin and eosin (H&E) and Verhoeff-Van Gieson (EVG) staining of coronary artery (CA) tissues from Kawasaki disease (KD) patients used in this study. The inset shows a magnification of the selected area. **e, f,** Microscope images showing H&E staining of control (CTRL) CA tissues used in this study. Abbreviations: CA, coronary artery; CTRL, control; E, epicardium; EVG, Verhoeff-van Gieson; KD, Kawasaki disease; LCx, left circumflex; M, myocardium; RCA, right coronary artery.

#### Extended Data Figure 2

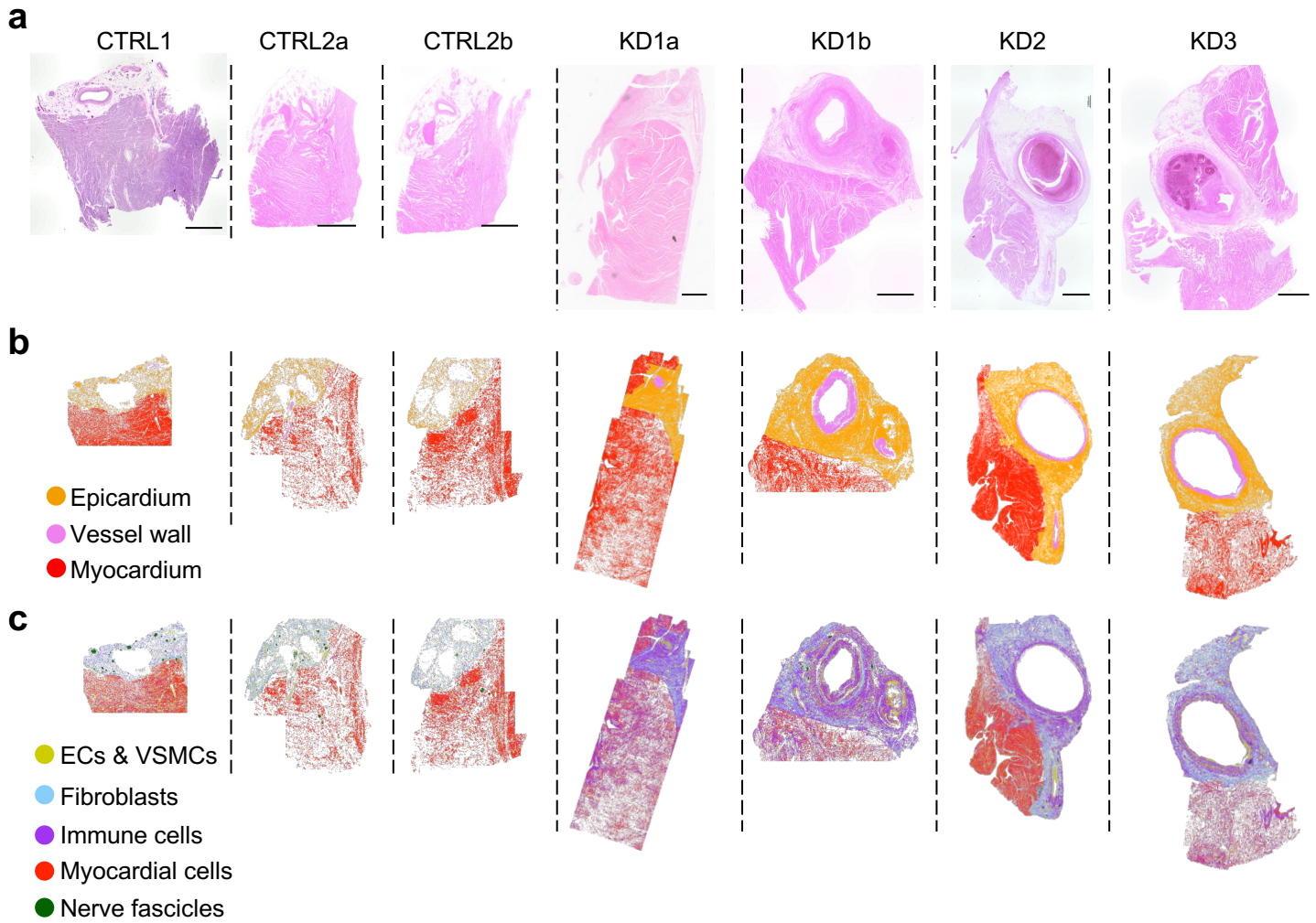

**Extended Data Figure 2. Multiplexed protein imaging resolves histological tissue layers in control and Kawasaki disease coronary artery tissues.** **a**, H&E staining of coronary artery (CA) tissues from control (CTRL) and patients with Kawasaki disease (KD) (as shown in Extended Data Fig. 1). Scale bars: 1 mm. **b**, Tissue layers (vessel wall, myocardium, and epicardium) identified based on the H&E and Napari imaging of slide topography. **c**, Spatial distribution of endothelial cells (ECs), vascular smooth muscle cells (VSMCs), fibroblasts, myocardial cells, and immune cells in each tissue section analyzed. Spatial identification of the different cell populations and tissue layers was achieved by differentially expressed canonical cell markers. Abbreviations: CTRL, control; ECs, endothelial cells; KD, Kawasaki disease; VSMCs, vascular smooth muscle cells.

Extended Data Figure 3

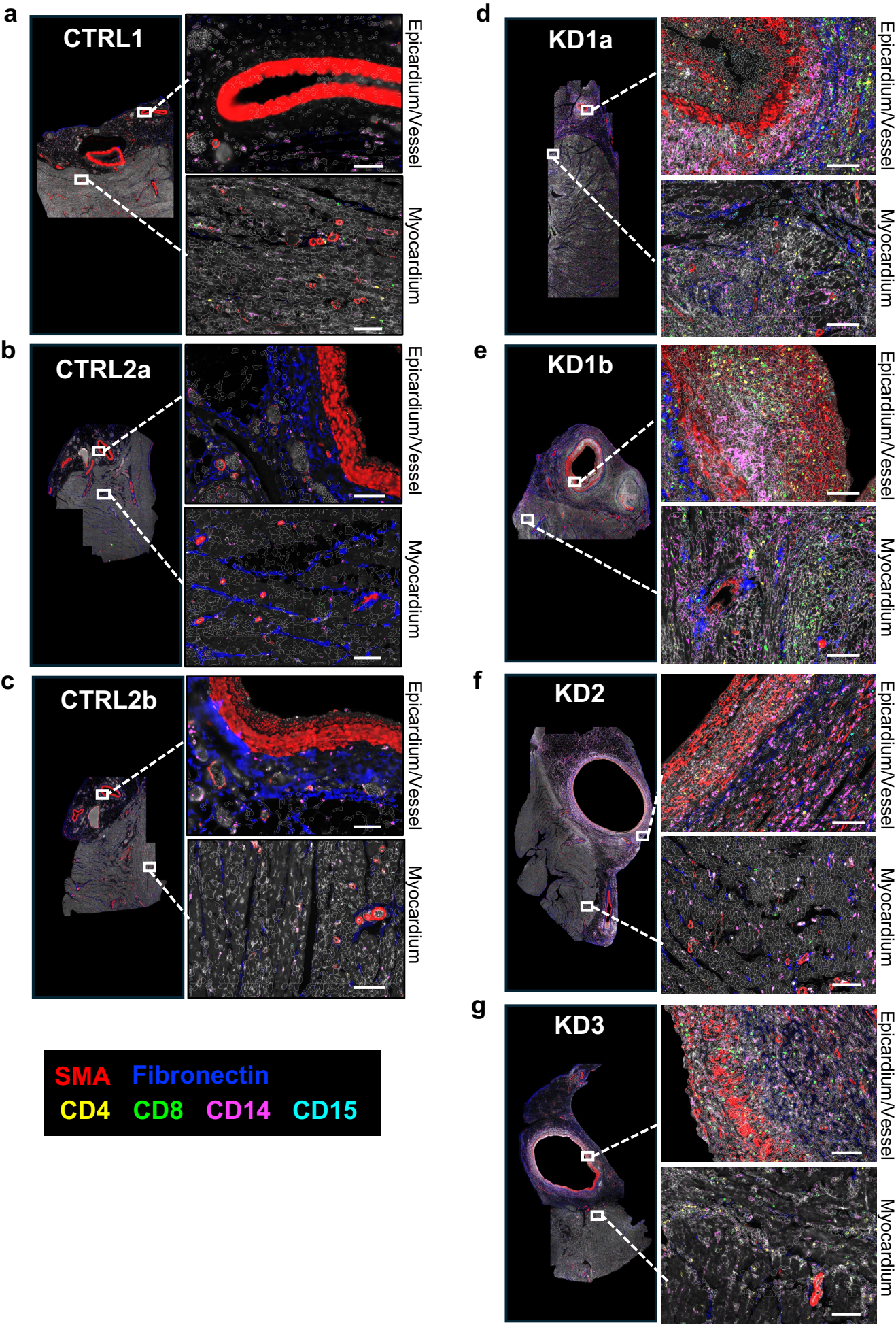

**Extended Data Figure 3. Cellular infiltrates in coronary artery tissues from patients with Kawasaki disease.** **a–g**, Napari images of coronary artery (CA) tissues and corresponding magnification images of the epicardium/tunica (top) and myocardium (bottom) from control (CTRL) and patients with Kawasaki disease (KD). Expression of the following proteins and associated cell types was assessed: SMA (smooth muscle cells), Fibronectin (fibroblasts), CD4 and CD8 (T cells), CD14 (macrophages), and CD15 (neutrophils). β2-macroglobulin (CD298/β2M) is shown in white. Scale bar: 100μm. Abbreviations: CTRL, control; KD, Kawasaki disease, SMA, smooth muscle alpha actin.

Extended Data Figure 4

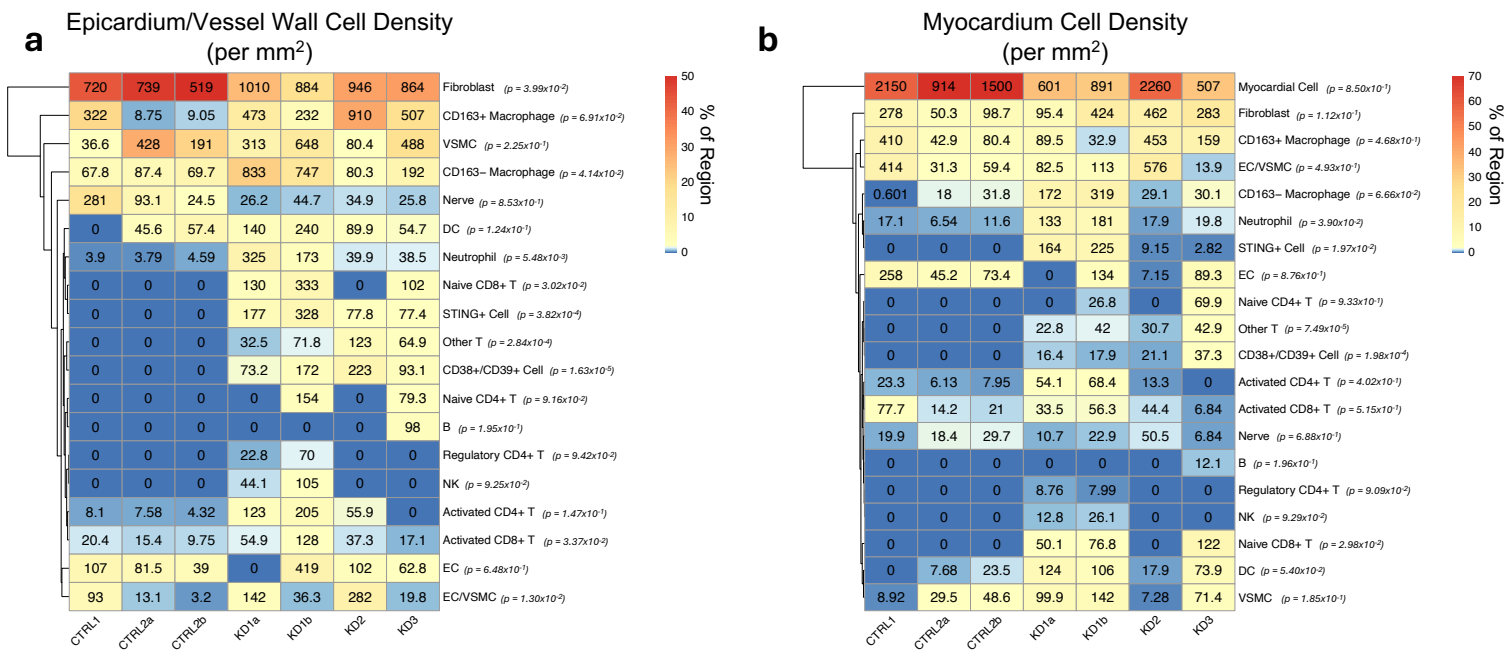

**c** Epicardium/Vessel Wall

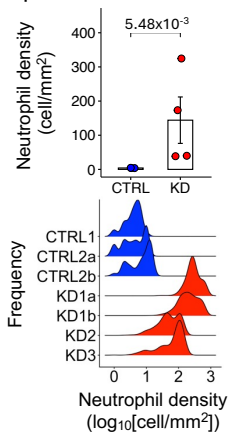

**d** CTRL1 CTRL2a CTRL2b

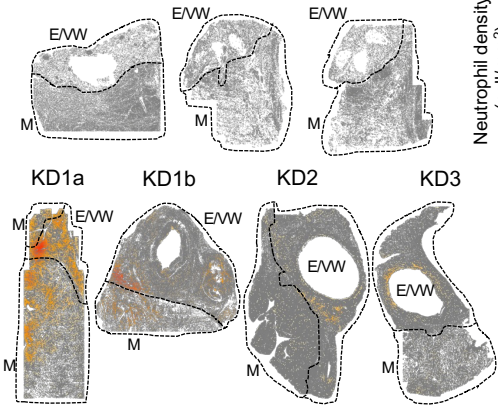

**e** Myocardium

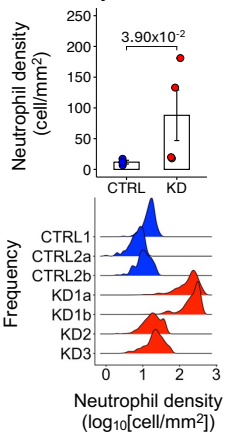

**f** Epicardium/Vessel Wall

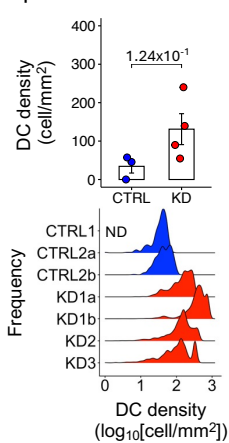

**g** CTRL1 CTRL2a CTRL2b

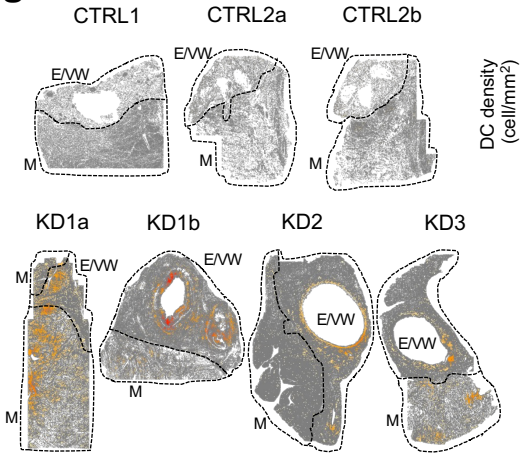

**h** Myocardium

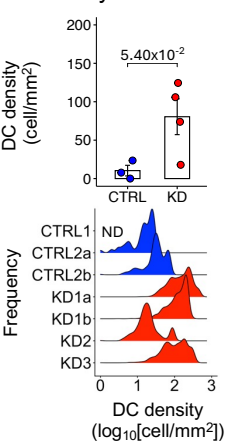

**Extended Data Figure 4. Immune cell densities in coronary artery tissues from control and patients with Kawasaki disease.** **a, b**, Heatmaps showing the density of all cell populations identified in the coronary artery (CA) epicardium and vessel wall (E/VW; **a**) and myocardium (M; **b**) of control (CTRL) individuals and patients with Kawasaki disease (KD). Cell density values (cell/mm<sup>2</sup>) are indicated within each heatmap cell. The color gradient (blue–yellow–red) represents the relative abundance (%) of each cell type for each tissue analyzed. *p*-value indicates statistical difference of the mean cell density in CTRL vs. KD samples. **c**, Bar plot and ridge plot of neutrophil density in the epicardium and CA vessel wall of CTRL vs. KD CA tissues. **d**, Spatial distribution of neutrophil density in the CA tissues from CTRL and KD patients. **e**, Bar plot and ridge plot of neutrophil density in the myocardium of CTRL and patients with KD. **f**, Bar plot and ridge plot of dendritic cells (DCs) density in the epicardium and vessel wall of CTRL and patients with KD. **g**, Spatial distribution of DCs density in CA tissues from CTRL and KD patients. **h**, Bar plot and ridge plot of DC density in the myocardium of CTRL and patients with KD. (**a-c**, **e**, **f**, and **h**) *p*-values calculated via the one-sided unpaired *t*-test. Abbreviations: CA, coronary artery; CTRL, control; E/VW, epicardium and vessel wall; KD, Kawasaki disease; M, myocardium; VSMC, vascular smooth muscle cells; EC, endothelial cells; DC, dendritic cells; NK, natural killer cells.

### Extended Data Figure 5

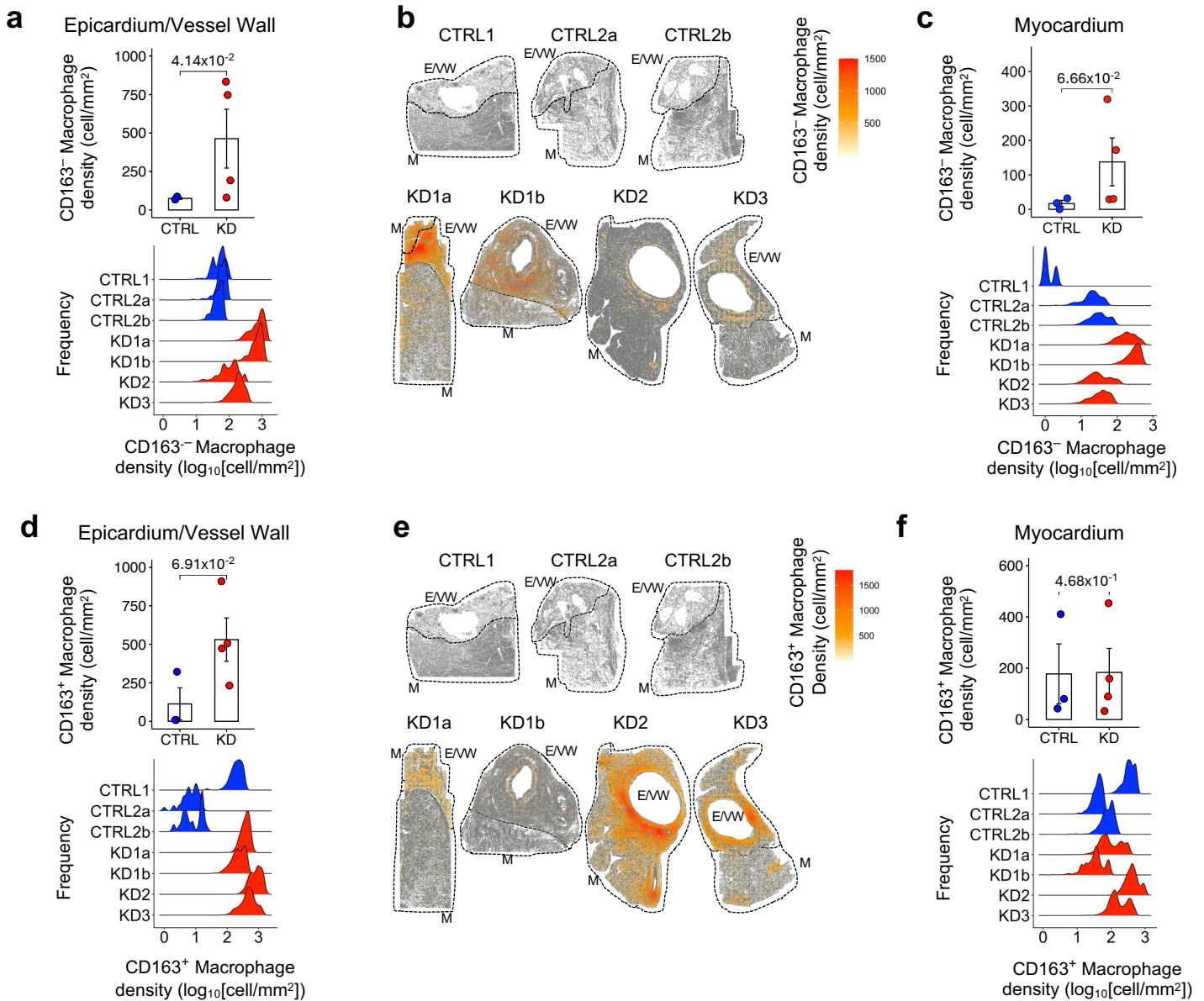

**Extended Data Figure 5. Densities of CD163<sup>-</sup> and CD163<sup>+</sup> macrophages in coronary artery tissues from control and patients with Kawasaki disease.** **a**, Bar plot and ridge plot of CD163<sup>-</sup> macrophage density in the coronary artery (CA) epicardium and vessel wall (E/VW) of control (CTRL) individuals and patients with Kawasaki disease (KD). **b**, Spatial distribution of CD163<sup>-</sup> macrophage density in CA tissues from CTRL and KD patients. **c**, Bar plot and ridge plot of CD163<sup>-</sup> macrophage density in the myocardium (M) of CTRL and KD patients. **d**, Bar plot and ridge plot of CD163<sup>+</sup> macrophage density in the epicardium and CA vessel wall of control and KD tissues. **e**, Spatial distribution density of CD163<sup>+</sup> macrophages in CA tissues from CTRL and KD patients. **f**, Bar plot and ridge plot of CD163<sup>+</sup> macrophages in the myocardium of CTRL and KD tissues. **a**, **c**, **d**, and **f** *p*-values calculated via the one-sided unpaired *t*-test. Abbreviations: CTRL, control; E/VW, epicardium/vessel wall; KD, Kawasaki disease; M, myocardium.

### Extended Data Figure 6

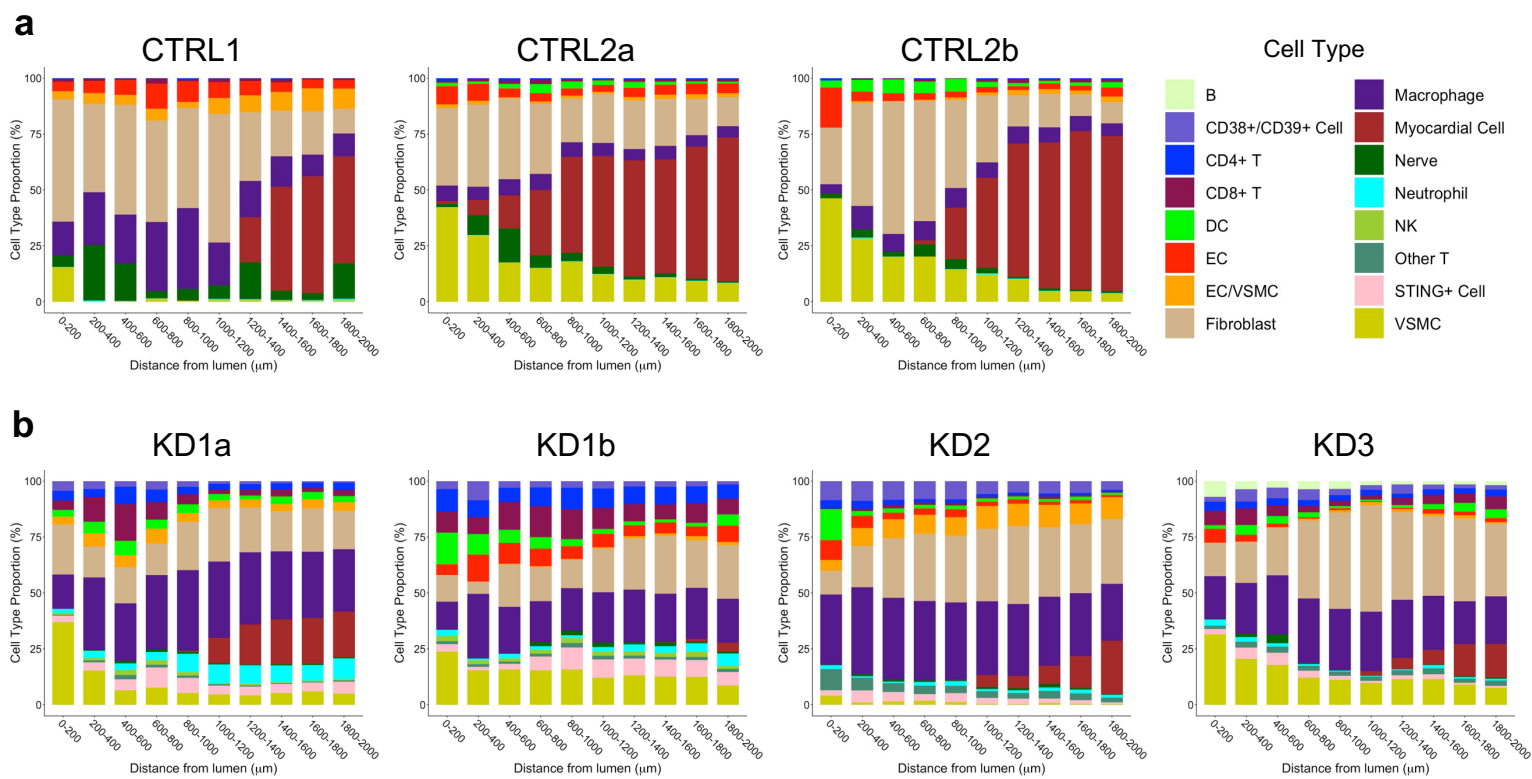

**Extended Data Figure 6. Lumen distances of immune cell infiltrations in coronary artery tissues from control and patients with Kawasaki disease.** **a**, Barplots showing cell type proportions at 200 μm intervals from 0 to 2 mm from the coronary artery (CA) lumen from control (CTRL) tissues. **b**, Barplots showing the cell type proportions at 200 μm intervals from 0 to 2 mm from the CA lumen from Kawasaki disease (KD) tissues. Abbreviations: CA, coronary artery; CTRL, control; KD, Kawasaki disease; VSMC, vascular smooth muscle cells; EC, endothelial cells; DC, dendritic cells; NK, natural killer cells.

Extended Data Figure 7

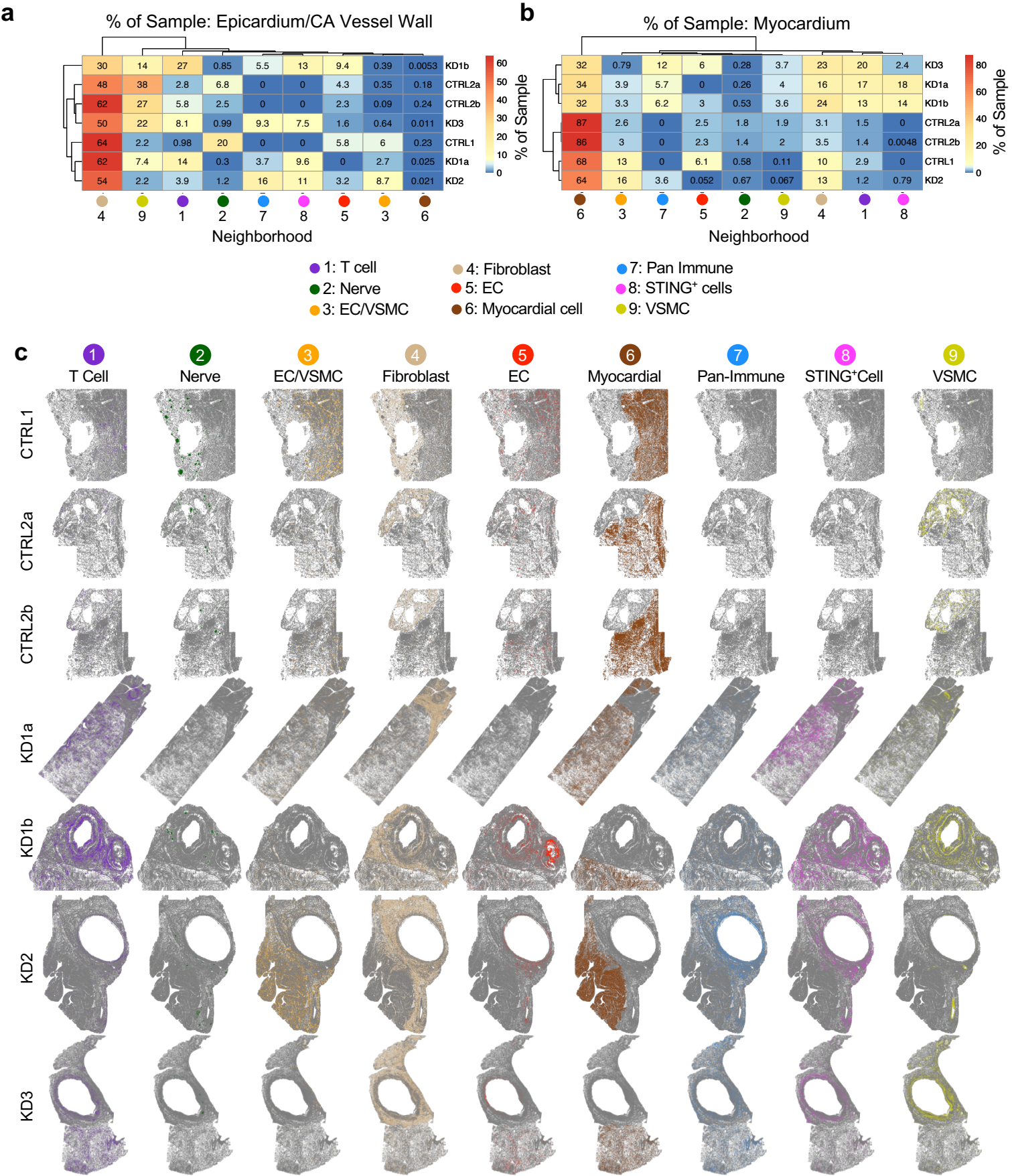

**Extended Data Figure 7. Neighborhood distribution in tissues from control individuals and patients with Kawasaki disease.** **a, b,** Heatmaps showing the distribution of the 9 identified neighborhoods in the epicardium/coronary artery (CA) vessel wall (a) and the myocardium (b) of control (CTRL) and patients with Kawasaki disease (KD). Values within each heatmap cell indicate the percentage of each neighborhood per sample. The color gradient (blue–yellow–red) indicates low to high percentage of each neighborhood. **c,** Spatial distribution of the identified 9 neighborhoods in each tissue section from CTRL and KD patients. Abbreviations: CTRL, control; KD, Kawasaki disease; VSMC, vascular smooth muscle cells; EC, endothelial cells; DC, dendritic cells; NK, natural killer cells.

### Extended Data Figure 8

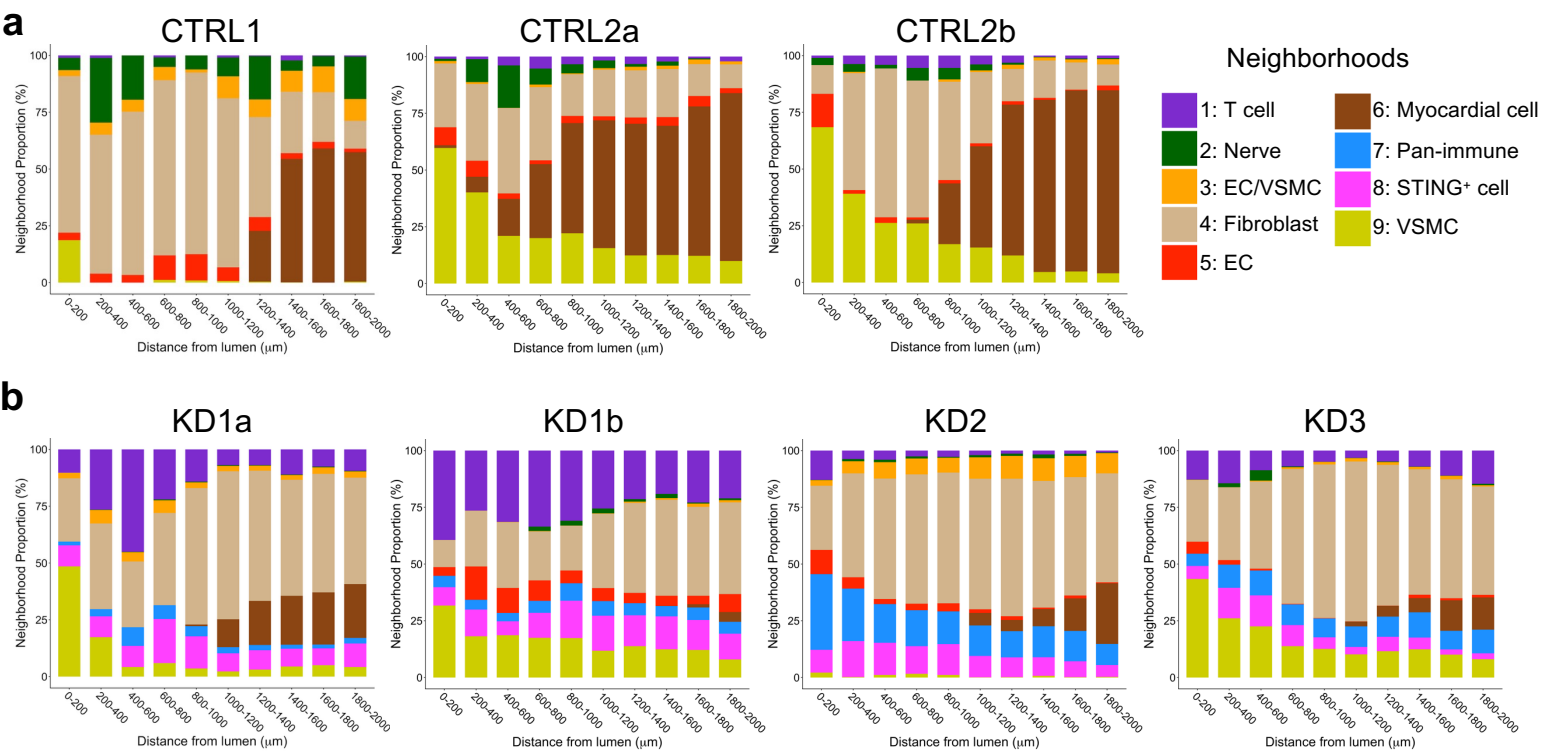

**Extended Data Figure 8. Coronary artery lumen distances of identified neighborhoods in tissues from control individuals and patients with Kawasaki disease.** **a**, Barplots showing neighborhood proportions at 200 μm intervals from 0 to 2 mm from the coronary artery (CA) lumen from control (CTRL) tissues. **b**, Barplots showing the neighborhood proportions at 200 μm intervals from 0 to 2 mm from the CA lumen from Kawasaki disease (KD) tissues. Abbreviations: CA, coronary artery; CTRL, control; KD, Kawasaki disease; VSMC, vascular smooth muscle cells; EC, endothelial cells.

#### Extended Data Figure 9

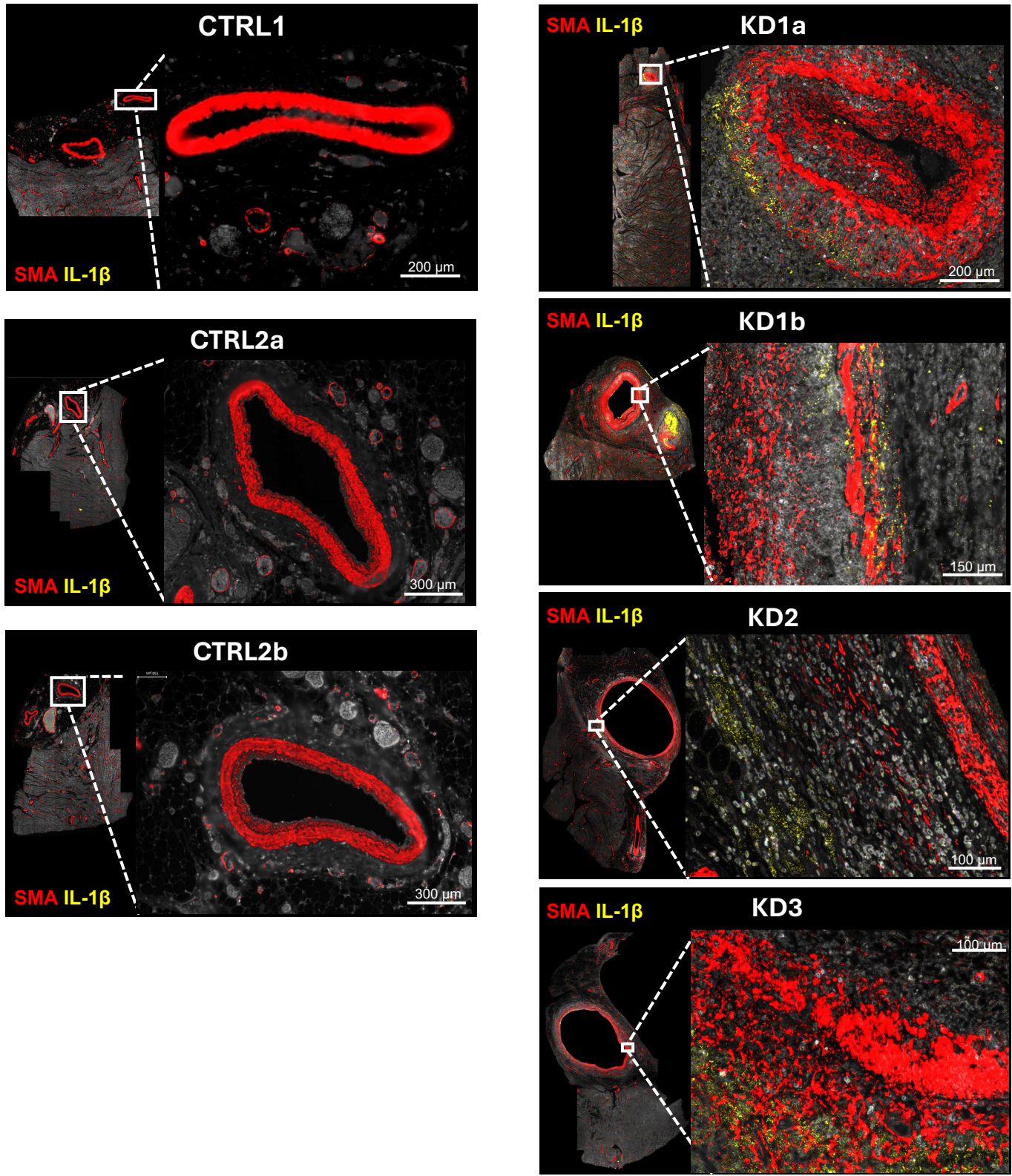

**Extended Data Figure 9. Detection of IL-1β in coronary artery tissues of patients with Kawasaki disease.** Fluorescence images of coronary artery (CA) tissues from control (CTRL1, CTRL2a and CTRL2b) individuals and patients with Kawasaki disease (KD1a, KD1b, KD2, KD3, KD4) showing the expression of SMA (red) and IL-1β (yellow). Images visualized using Napari. β2-macroglobulin (CD298/B2M) is shown in white. Abbreviations: CTRL, control; KD, Kawasaki disease; SMA, smooth muscle actin.

#### Extended Data Figure 10

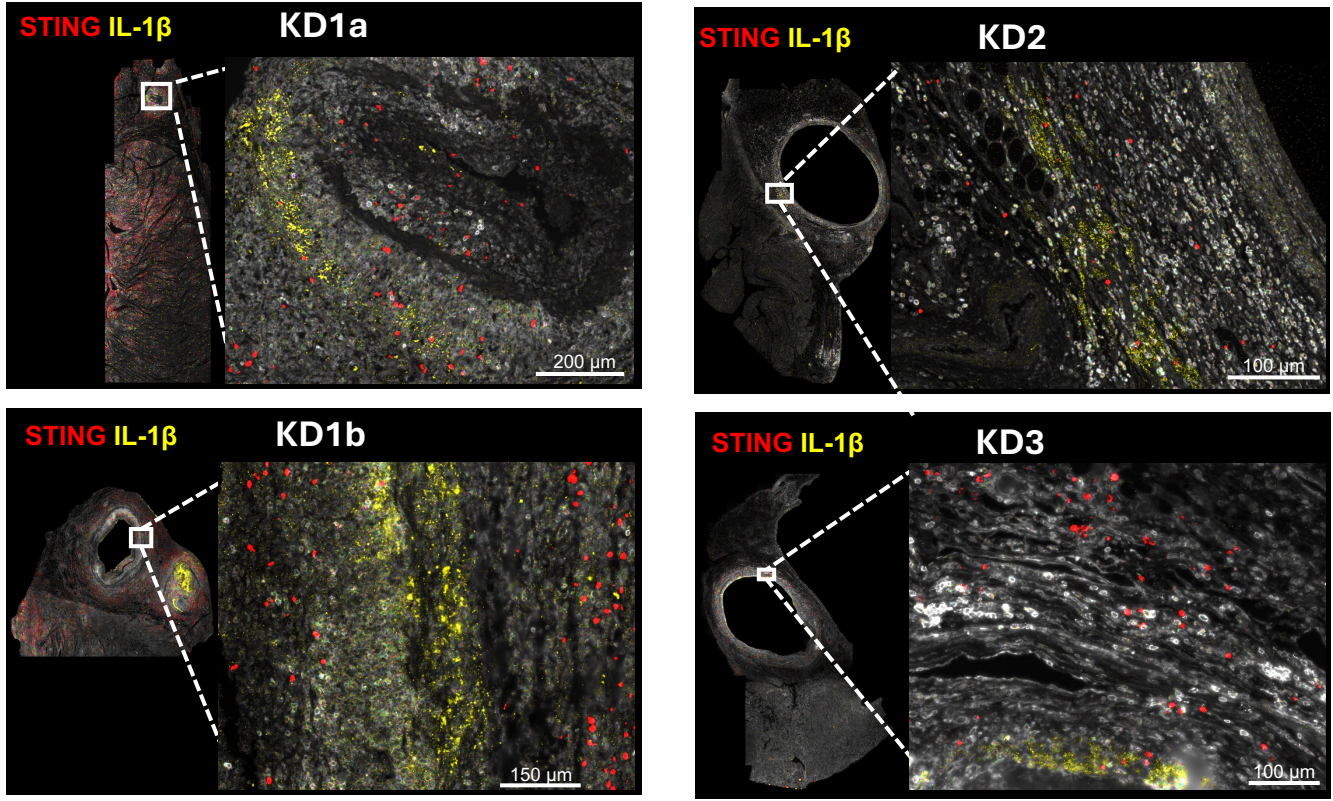

**Extended Data Figure 10. STING and IL-1β expression in coronary artery tissues from patients with Kawasaki disease.** Fluorescence images of coronary artery (CA) tissues from patients with Kawasaki disease showing the expression of STING (red) and IL-1β (yellow). β2-macroglobulin (CD298/B2M) is shown in white. Images visualized using Napari. Abbreviations: KD, Kawasaki disease.

Extended Data Figure 11

a

GSE68004

Acute KD (n=76) vs. healthy control (HC; n=37)

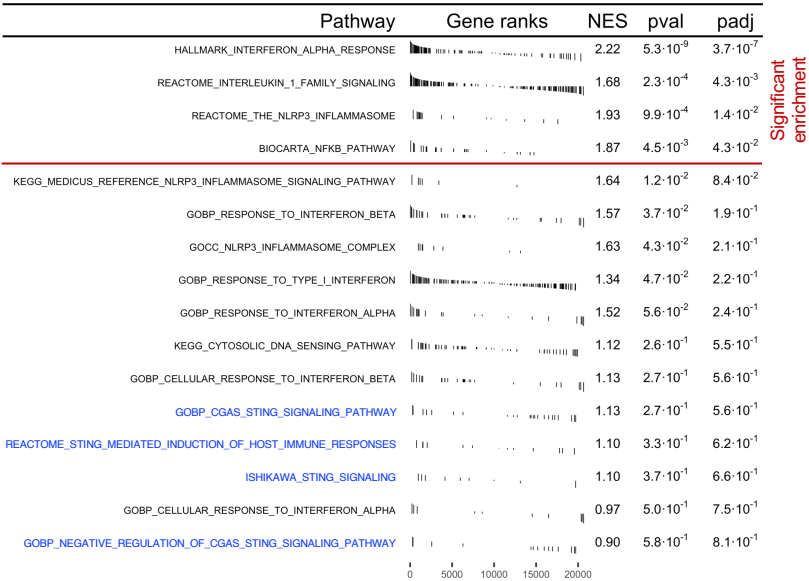

Significant enrichment

b

GSE178491

Acute KD (n=105) vs. febrile control (FC; n=30)

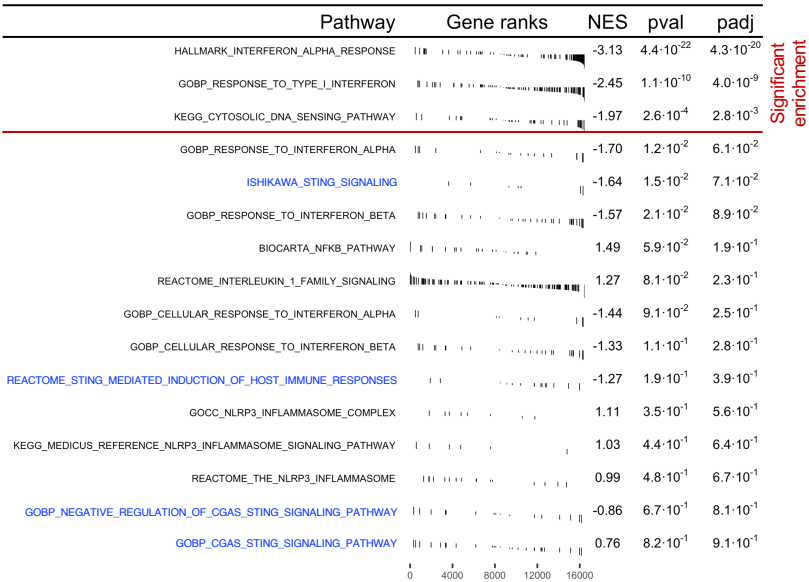

Significant enrichment

c

GSE178491

Acute KD (n=105) vs. convalescent KD (n=31)

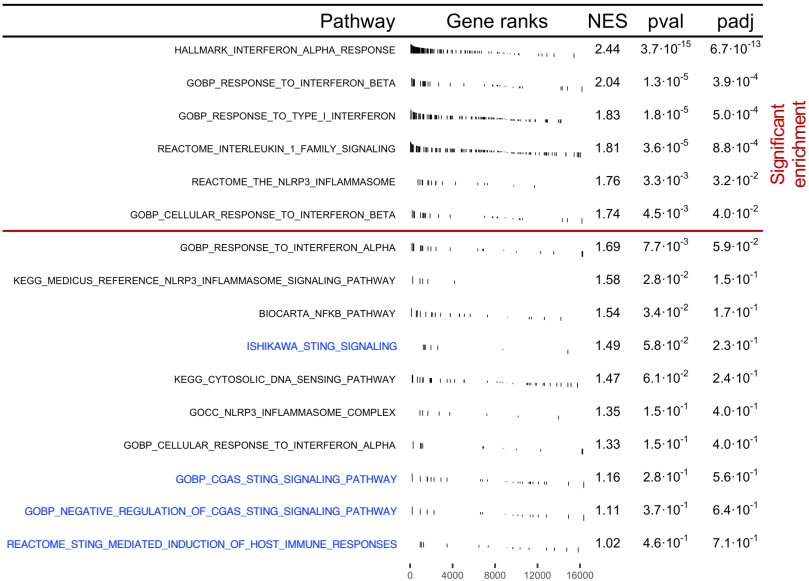

Significant enrichment

**Extended Data Figure 11. Gene set enrichment analysis (GSEA) of whole blood transcriptomics datasets of control individuals and patients with Kawasaki disease.**

Publicly available gene expression datasets generated from the whole blood of healthy controls (HC), febrile controls (FC), patients with acute Kawasaki disease (KD), and convalescent KD patients (GSE68004 and GSE178491) were analyzed by GSEA. **a**, Enrichment of pathways in whole blood samples of patients with acute KD (n = 76) vs. HCs (n = 37) (GSE68004). **b**, Enrichment of pathways in whole from patients with acute KD (n = 105) vs. FCs (n = 30) (GSE178491). **c**, Enrichment of pathways in whole blood from acute KD patients (n = 105) vs. convalescent KD patients (n = 31) (GSE178491). Red lines separate significant pathways (*adj. p-value* < 0.05) from those that are not significantly enriched. Blue text indicates cGAS-STING gene-related pathways. Abbreviations: HC, healthy control; FC, febrile control; KD, Kawasaki disease.

Extended Data Figure 12

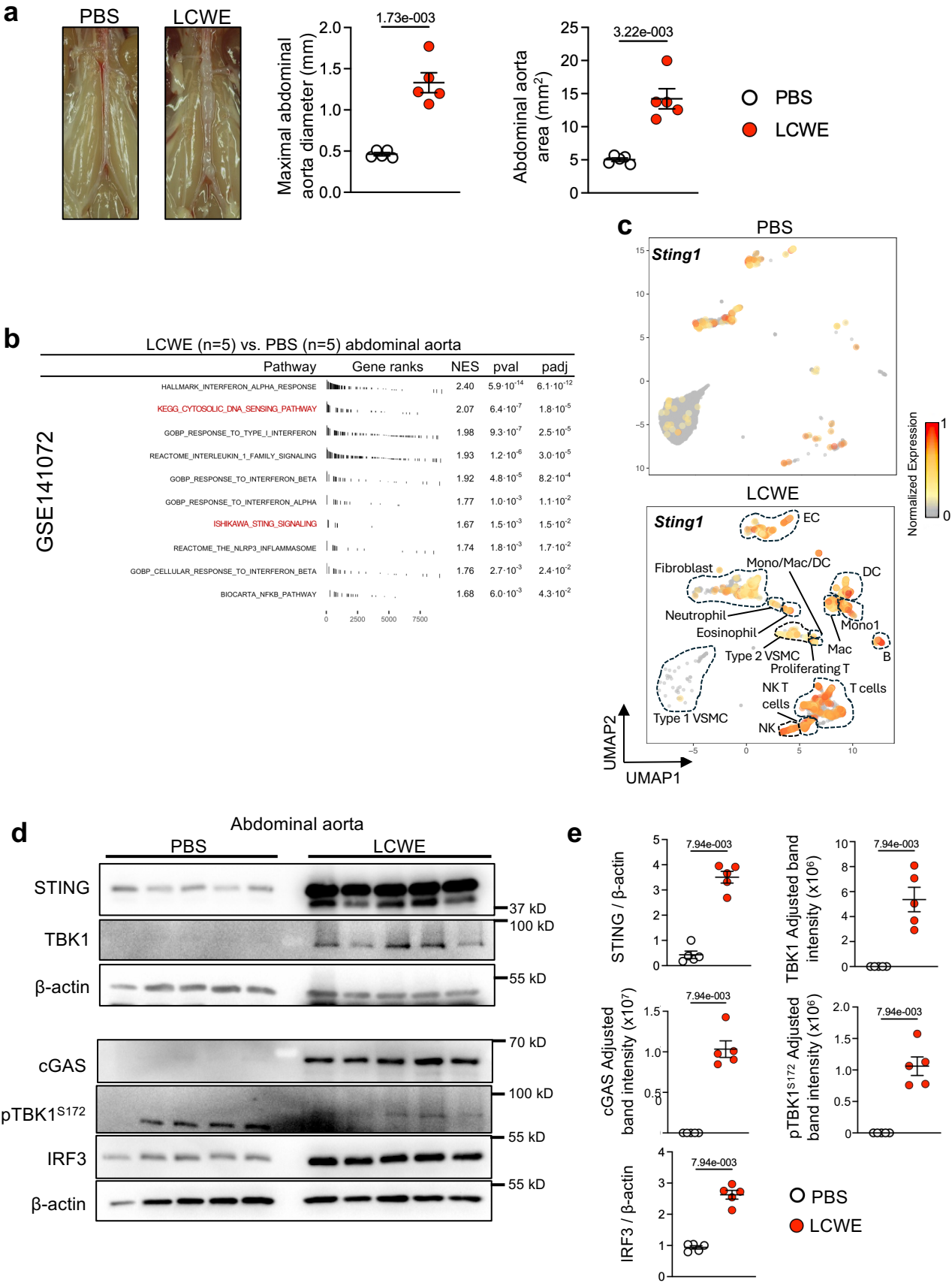

**Extended Data Figure 12. Activation of the cGAS-STING pathway in abdominal tissues from LCWE-injected mice.** **a**, Pictures of the abdominal area, maximal abdominal aorta diameter, and abdominal aorta area measurements of PBS- and LCWE-injected WT mice (n = 5/group). **b**, Gene set enrichment analysis (GSEA) of significantly enriched (*adj. p-value* <0.05) selected gene sets related to cGAS-STING signaling, inflammatory responses, and type I interferon response (**Supplementary Table 3**) in a published bulk RNA sequencing dataset generated from abdominal aortas of PBS- and LCWE-injected mice (GSE141072; n = 5/group). All genes were ranked by log<sub>2</sub> fold change (FC) between LCWE vs. PBS. Vertical marks indicate the position of each gene within the ranked list. Normalized enrichment scores (NES) indicate enrichment of the pathway in LCWE-injected abdominal aortas. Blue text indicates cGAS-STING gene-related pathways. **c**, UMAP plot of cells expressing *Sting1* in abdominal aorta cells isolated from PBS (n = 5386 cells) and LCWE-injected (n = 5091 cells) WT mice. (published dataset GSE178765). **d**, Western blot (WB) analysis of STING, TBK1, cGAS, pTBK1<sup>S172</sup> and IRF3 in abdominal aorta lysates from PBS- and LCWE-injected WT mice at 2 weeks post-injection (n = 5/group). **e**, Quantification of protein expression by densitometry in abdominal aorta tissue lysates from PBS- and LCWE-injected WT mice, at 2 weeks post-injection. STING and IRF3 expression were normalized to β-actin and are presented as FC. cGAS, TBK1, and pTBK1<sup>S172</sup> are presented as adjusted band intensities, where PBS samples with no detectable band after background subtraction were assigned a value of “0”. Data are presented as mean ± SEM (a, e), with each symbol representing one individual mouse. *p-values* obtained using the Mann-Whitney test (a, e).
